# Lipid droplet-sphingolipid crosstalk regulates Doa1 activity and DNA damage response

**DOI:** 10.64898/2025.12.12.694079

**Authors:** Ranu Singh, Raghuvir Singh Tomar

## Abstract

Neutral lipid droplets (nLDs) are dynamic storage organelles that protect the cells from lipotoxicity. They store excess neutral lipids, sequester heavy metals and serve as membrane reservoirs. However, their role in genome integrity remains unclear. Previous studies have linked the shortened lifespan of LD deficient cells to impaired mitochondrial functions caused by defective sterol metabolism, but the molecular origin of this dysfunction remains elusive. In this study, we demonstrate that lack of nLDs inhibits DNA damage sensing mechanism, causing insensitivity to DNA damaging agents. Despite sustained DNA breaks, LD deficient cells unable to activate the checkpoint kinases or arrest cell cycle progression. Persistent insensitivity to double strand breaks results in accumulation of mutations, loss of nuclear as well as mitochondrial DNA integrity leading to accelerated cellular ageing. Through systematic genetic dissection of nLD synthesis genes, we identify a crucial role of lanosterol esterification in lipid droplet formation and their role in the regulation of DNA damage sensing and repair responses. Our study identifies a key regulatory protein, Doa1 that functionally links lipid droplet biosynthesis to DNA damage repair kinase signalling. Together, these findings establish a previously uncharacterized role of neutral lipid droplets in the maintaining the genome stability by acting not only as metabolic buffers but also as signalling platforms that coordinate DNA damage sensing and repair processes. Our work reveals a new mechanistic connection between cellular lipid homeostasis and genome maintenance, highlighting lipid droplets as critical determinants of healthy cellular aging.

## Introduction

Lipids are fundamental to cellular homeostasis, which serve as dense energy stores, essential components of membrane and secondary messengers in signalling pathways. Their neutralization into lipid droplets is crucial to sequester potentially damaging free fatty acids and other lipid species (1).

Lipid droplets/Neutral lipid droplets (nLDs) are dynamic, mono-layered storage organelles that originates from endoplasmic reticulum (ER) membrane. Their primary storage components are triacylglycerol (TAG) and sterol esters (SE) which acts as transient energy reservoir and buffers against lipotoxic stress. In *Saccharomyces cerevisiae,* TAG synthesis is catalysed by diacylglycerol acyl-transferase encoded by dga1 and lro1 and sterol ester formation is catalysed by acyl-coenzyme A (CoA): cholesterol acyl transferases (ACATs), Are1 and Are2 (2). Among these, Are1 preferentially esterifies lanosterol while Are2 acts mainly on ergosterol (3). Yeast strain lacking these four genes fails to form lipid droplet and show significant alteration in their TAG and sterol synthesis (4). Thus, nLD formation is essential for the neutralisation and sequestration of excess lipids, protecting the cell from lipotoxicity.

Beyond their canonical role in lipid storage, LDs have emerged as multifunctional organelles that participate in diverse cellular processes related to stress adaptation and organelle communication (5–7). Under the nutrient starvation conditions, lipids released through autophagy are rapidly esterified to TAGs and stored in lipid droplets (8). Lipid droplets also contribute to the cytokinesis by supplying components necessary for proper septin assembly and acts as membrane reservoir in the cells (9). Studies suggest that in the nucleus, lipid droplets regulate phospholipid synthesis by modulating intranuclear interactions between two rate limiting factors OPI1 and CCTα on their surface (6). Excess histones get stored into the lipid droplets to prevent their degradation during *Drosophila* oogenesis (10). Lipid droplets have been shown to sequester toxic compounds and facilitates their exocytosis highlighting their protective role under adverse environmental stress conditions (11,12)

Upon genotoxic exposure, cells accumulate lipid droplets in dose dependent manner. Evidence suggests that the ATM/ATR axis of DDR pathways function as a key mediator linking lipid droplet biogenesis to DNA damage stress. For example, in *C. elegans* the lipid droplets utilizes Mafr-1 (a negative regulator of RNA Pol III) along with atm-1 (homologe of Tel1) and atr-1 (homolog of Mec1) kinases of DDR pathway in response to harmful UV exposue (13). In yeast as well as in mammalian RPE cells lines, the selective storage of sterols in lipid droplets has been reported to bind and attenuate the DDR kinase, ATM through Phosphoatidyl-inositol-4-phosphate. This attenuation of DDR kinase mitigates the signalling pathways of DNA damage and repair processes (14).

The DNA damage response (DDR) represents a highly conserved signaling network that maintains genomic integrity by detecting, signalling and repairing DNA lesions. In yeast cells, two central kinases, Tel1 and Mec1, homologs of mammalian ATM and ATR, respectively, serves as primary sensors of DSBs (DNA-double strand break) and replication stress. Tel1 is recruited to DSB through MRX complex (Mre11-Rad50-Xrs2), while Mec1 associates with Ddc2 and binds to single-stranded DNA lesions coated by RPA (ssDNA binding protein) (15,16). After activation, both kinases phosphorylates downstream effectors mainly Rad53 and Chk1, initiating check point signalling.

Activated Rad53 stabilizes replication fork, and enhances DNA repair by increasing dNTP synthesis through Sml1 degradation, and transcriptional de-repression of *RNR* genes. Chk1, in parallel, stabilizes the anaphase inhibitor Pds1, enforcing cell-cycle arrest untill DNA integrity is restored (17). Collectively these pathways coordinates replication, repair and checkpoint regulation essential for the genome stability under genotoxic stress conditions.

Previous studies have reported a positive correlation between Rad53 phosphorylation and number of intracellular lipid droplet molecules during genotoxic stress conditions. An increase in Rad53 activation often coinsides with elevated lipid droplet formation, suggesting two possible interpretations, lipid droplets may either function as upstream sensors that modulate DNA damage response or downstream regulators of Rad53-mediated signaling. However, despite this consistent parallel infomations about the role of lipid droplets, no direct mechanistic link has been established between lipid droplet biogenesis and DDR signaling. The concurrent increase in both, lipid droplet biogenesis and DDR signaling under DNA damage condition, strongly implies a functional connection between these two pathways, however, the precise nature of this relationship remains unclear. Considering the persistence and abundance of lipid droplets following DNA damage, it is plausible that they contributes to cellular protection and DNA repair processes. Therefore, in this study, we examined DNA damage sensing and repair responses in the absence of lipid droplet biosynthesis. We studied the response of the lipid droplet deficient mutants on different DNA damaging agents. We observed lipid droplet-deficient cells to be hypersensitive to hydroxyurea, but resistant to direct DNA damaging agent, methymethanesulfonate and UV. Further biochemical analysis revealed that this unexpected resistant growth phenotype of lipid droplets deficient mutant cells to MMS is due to the dysregulation in DDR sensing, characterized by low Rad53 phosphorylation. This sensing defect arises from elevated Doa1 expression and non-specific ubiquitination of cellular proteins in lipid droplet deficient mutant. We further show that the lipid droplet deficient mutant has increased pool of sphingolipd that gets neutralized and stored by droplet formation in the wild type cells. This active sphingolipid pool triggers the Doa1 hyperactivity. The cumulative effect of this dysregulation leads to defect in DNA damage checkpoint signalling which results in accumulation of DNA damage and decreased chronological life-span. Finally, the experiments with combinatorial deletion mutats of LD biosynthesis genes revealed the regulatory role of Are1 in co-ordinating both lipid droplets biosynthesis and Rad53 phosphorylation. Together our finding uncovers a regulatory axis connecting lipid droplet homeostasis to DNA damage sensing machinery. Understanding this relationship provide new insights into potential regulatory role of lipid droplets in genome survillience and cellular stress adaptation.

## Results

### Lipid droplet deficiency confers general stress sensitivity but unexpected resistance to direct DNA-damaging conditions

To understand the role of lipid droplets under genotoxic stress conditions, we utilized quadruple deletion mutant where all four genes related to TAG and sterol ester synthesis are deleted. We compared the growth of wild type (SCY62) and lipid droplet deficient mutant, H1246; a quadruple gene deletion mutant; are1Δ/are2Δ/dga1Δ/lro1Δ, upon treatment with hydroxyurea (HU), methyl methane sulphonate (MMS), and ultraviolet (UV) radiation. Hydroxyurea depletes dNTP pool without directly affecting DNA integrity, whereas MMS introduces alkylating lesions at N7-guanine and N3-adenine sites, generating apurinic site and strand breaks during base excision repair (18). On the other hand, UV irradiation induces cyclobutene pyrimidine dimers between adjacent thymine or cytosine bases.

Since the previous studies suggest a protective role of lipid droplets under genotoxic conditions, we hypothesized that in their absence, cells will become hypersensitive to DNA damaging agents. To examine this hypothesis, we spotted the cells on SC agar plates containing 150mM, and 200mM of HU, 0.01%, and 0.015% of MMS (Figure 1A, S1Ai-Aii). To examine the effect of UV radiation, different dilutions of cells were first spotted on SC agar plate and then exposed with 6×10^4^µJ/cm^2^ radiation in UV cross linker. Spot assay analysis revealed hypersensitivity of LD deficient mutants (H1246 and Q47) to hydroxyurea treatment. However, on the other hand, the same mutant exhibited markedly improved survival on MMS and UV treatments relative to wild type cells. Growth-curve analysis in liquid SC growth media in presence of MMS also showed the same growth kinetics. Although the H1246 quadruple lipid droplet deficient mutant grows slower than wild type in untreated condition, it consistently displayed better survival under MMS treatments (Figure 1B).

**Figure 1:**
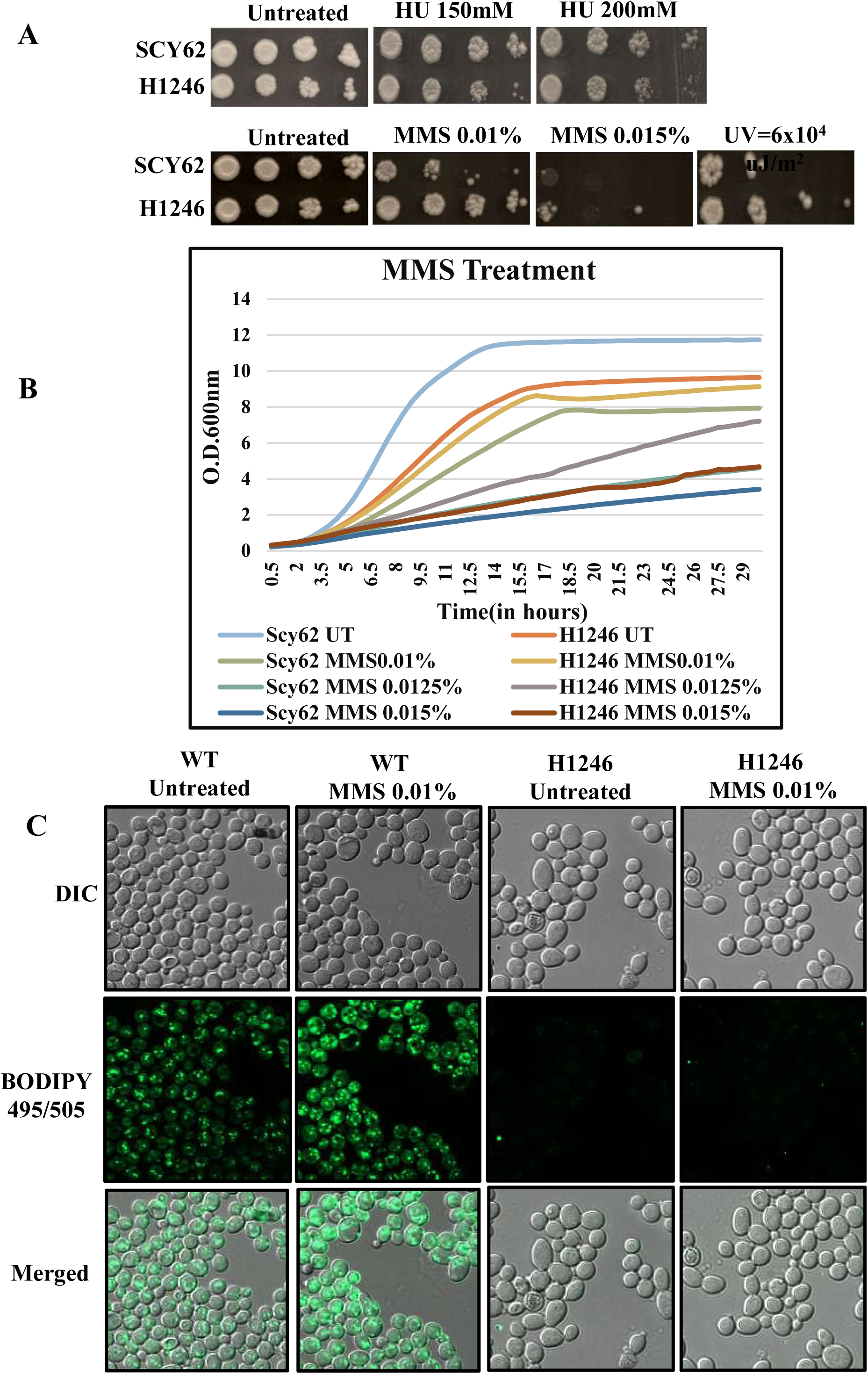
Lipid droplet mutant shows resistance towards MMS and UV treatment. **(A)** Spot assay of wild type (SCY62) and lipid droplet deficient mutant (H1246) were grown on plates containing methyl methane sulphonate (MMS), hydroxyurea (HU) and ultra-violet (UV) irradiation. Tenfold-serial dilutions were spotted onto the synthetic complete media agar plate containing 0.01%, and 0.015% of MMS, 150mM and 200mM of HU along with untreated control plate. UV irradiation was performed after spotting the cells on SC agar media. Growth was monitored after incubation at 30°C for 2-4 days and scanned using HP scanner. **(B)** Growth curve analysis of SCY62 and H1246 cells grown in liquid media containing increasing concentrations of MMS. Optical density (OD_600_) was recorded at 30min interval in Bio-Tek plate reader for 30 hours to assess effect of MMS on the growth. **(C)** Fluorescence microscopy of cells stained with BODIPY (493/503nm) dye to measure intracellular levels of lipid droplets. SCY62 and H1246 cells were stained following 0.01% MMS treatment and images were taken by Zeiss-apotome microscope to compare lipid droplet abundance between strains. Experiments were performed a minimum of three times (three independent biological replicates), and the representative image is shown here.

Because several of the DNA damaging agents such as camptothecin, zeocin, ionizing radiation are known to increase LD formation, we used BODIPY (493/503) dye to examine the LD count under MMS treatment condition (19). BODIPY fluorescence suggest significant increase in the number of lipid droplets in wild type cells following 0.01% MMS treatment, whereas H1246, LD deficient mutant as expected remained completely devoid of LDs as it did not show any florescence (Figure 1C, S1C-E).

Among the four LD biosynthesis genes, *ARE1* esterifies lanosterol and *ARE2* esterifies ergosterol. Any defect in lanosterol or ergosterol esterification dysregulate the cytosolic ergosterol pool. Ergosterol plays an essential role to maintain the plasma membrane integrity. Cells lacking ergosterol show compromised membrane integrity and permeability (20). Since lipid droplet deficient mutant also lacks these two genes, it is possible that membrane permeability of this mutant is compromised, which might affect the MMS intake. To determine whether the MMS tolerance observed in the quadruple mutant resulted from a global membrane defect or is it an altered DDR, we examined their growth under other stress inducing agents including cell wall perturbing agent (CFW and CR), osmotic stress (NaCl), oxidative stress (paraquat, menadione), unsaturated and saturated fatty acids (palmitoleic and oleic acid, stearic acid respectively), heavy metals and high temperature stress (37°C). Membrane permeability plays decisive role in intake and tolerance of these stress inducers. We spotted the cells on calcofluor white (CFW, 25 and 50µg/ml) Congo-red (CR, 50 and 100µg/ml), NaCl (1M), paraquat (0.5mM and 1mM), menadione (0.1mM and 0.2mM), ethanol (2%), glycerol (2%), acetic acid (0.1% and 0.2%), stearic acid (0.2%), palmitoleic acid (0.1% and 0.2%), oleic acid (0.2%), Rapamycin (10ng/ml), FeSO_4_ (10Mm), CuSO_4_ (0.5mM), CdCl_2_ (5mM), AlCl_3_ (2mM), MnSO_4_ (7.5mM), ZnSO_4_ (5mM), AsCl_3_ (7.5µM) and 37°C (Figure S1F-G). Despite their slow basal growth, the lipid droplet deficient mutant (H1246) showed growth phenotype same as wild type cells or a very mild growth defect in above treatment conditions. These observations indicate that membrane permeability of the lipid droplet deficient mutant is not very different that the wild type cells. Importantly, the growth of nLD deficient mutant (H1246) was found to be almost same as that of wild type cells in all the above treatments except Rapamycin. The mutant, H1246 showed resistant growth phenotype under rapamycin treatment condition which need to be explored further (Figure S1G).

Collectively, these finding suggest that resistance growth phenotype of lipid droplet deficient mutants to MMS/UV treatments is not due to altered membrane permeability or non-specific adaptation. Instead, it reflects a paradoxical and specific defect associated to DNA damage sensing and downstream signal processing.

### nLD mutants exhibit reduced DDR kinase activation

Given the unexpected resistant growth phenotype of H1246 mutant to MMS and UV treatments, we sought to determine whether this phenotype reflects alteration in DNA damage sensing or the repair process. To test this possibility, we first examined the activation state of DDR effector kinase, Rad53. Wild type and H1246 lipid droplet deficient mutant cells were exposed to increasing concentrations of MMS (0.01%, 0.0125% and 0.015%) for two hours, and whole cell extracts were prepared to measure the Rad53 phosphorylation by immunoblotting. As expected, the wild type cells exhibited progressive increase in Rad53 phosphorylation, which is indicative of normal checkpoint activation (Figure 2A, S2A-C). In contrast, LD deficient mutant cells displayed markedly reduced Rad53 phosphorylation upon MMS treatments, indicating a profound defect in DNA damage checkpoint activity.

**Figure 2:**
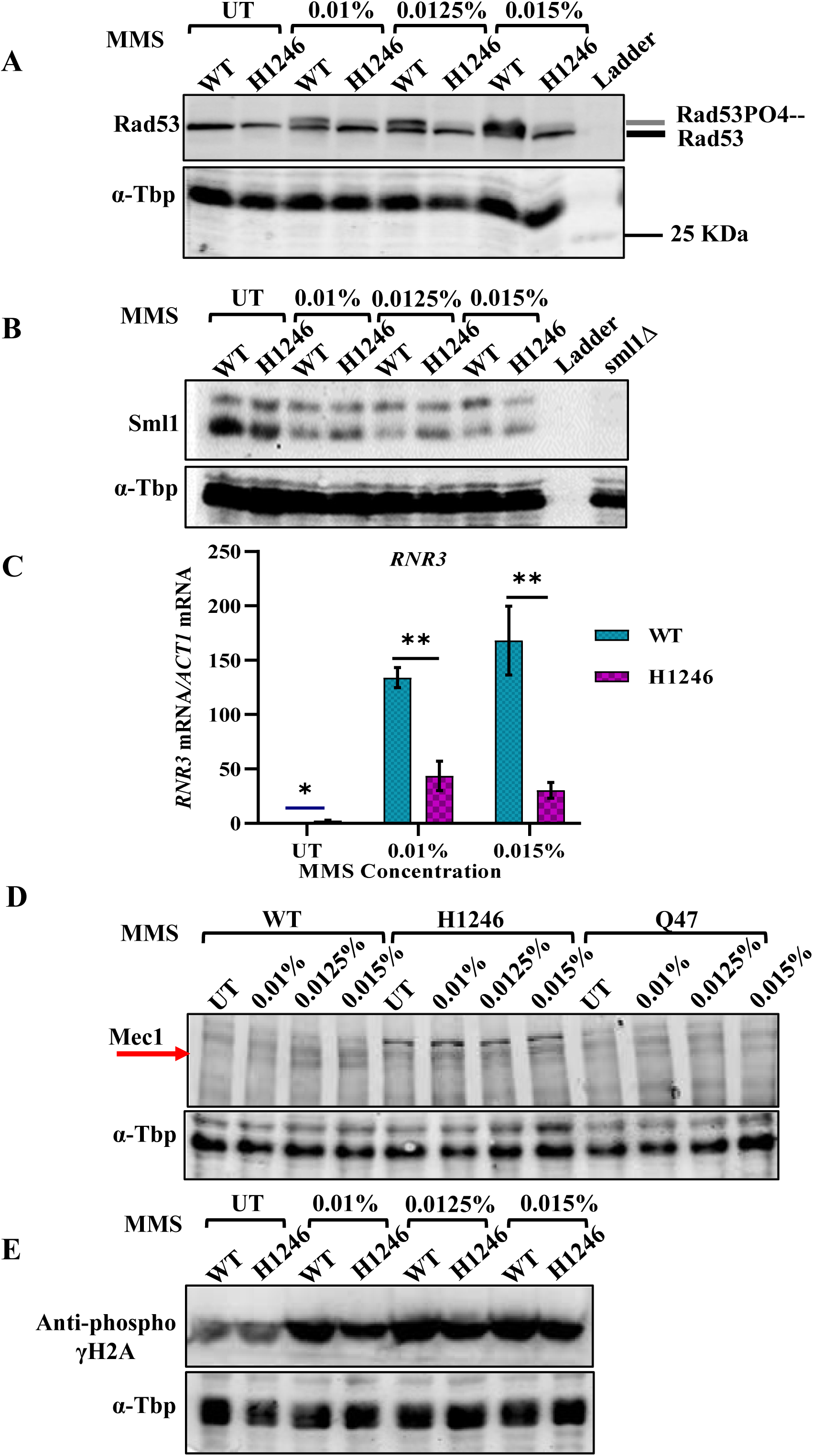
Lipid droplets deficiency impairs with Rad53 phosphorylation, Sml1 degradation and *RNR* induction. **(A)** Wild type and H1246 mutant cells were treated with increasing concentrations of MMS (0.01%, 0.0125%, 0.015%) for 2 hours and whole cell extracts were prepared using TCA precipitation method. Rad53 phosphorylation was assessed by western blotting, blots were developed with Bio-Rad ECL kit. α-TBP antibody was taken as a protein loading control. H1246 mutant showed significantly reduced Rad53 phosphorylation across all the MMS treatments. **(B)** Under similar MMS treatments as in panel A, Sml1 protein degradation were examined by western blotting to monitor the signalling downstream of Rad53 phosphorylation. Sml1Δ strain was taken as negative control and western blotting using α-Tbp antibody was performed for the protein loading control. **(C)** RT-qPCR analysis to measure the expression of *RNR3* in wild type and H1246 mutant cells in untreated and upon MMS treatments. The mRNA expression level of *RNR3* relative to *ACT1* mRNA was measured. Fold change was calculated using the formula 2*^-^*^ΔΔ*CT*^, where ΔΔ*CT* is Δ*CT* (test gene)*-*Δ*CT* (control gene). The data represents the mean from three independent biological replicates with the bar depicting SEM (standard error mean). Student’s t-test was performed to calculate statistical significance. **(D-E)** Western blotting to measure total Mec1 phosphorylation and γ-H2A phosphorylation in wild type, H1246 and Q47 mutants in untreated and MMS treatments. Experiments were performed a minimum of three times (three independent biological replicates), and only representative image is shown here.

The phosphorylation of Rad53 is catalysed by Mec1/Tel1 (homologue of ATR/ATM) kinase which in turn phosphorylates Dun1. Phosphorylated Dun1 removes the transcription repressor Crt1 from Rnr genes (*RNR2, RNR3* and *RNR4*) and allow their expression, which contributes to dNTP pool required for efficient DNA repair. Simultaneously, Rad53 also phosphorylates Sml1, which is the repressor of Rnr1 protein complex protein. Phosphorylated Sml1 protein undergoes degradation that leads to increase in dNTP pool (21). Given that Rad53PO^4--^ triggers the degradation of Rnr1 inhibitor, Sml1, we next compared the Sml1 protein levels under MMS treatment conditions in wild type and nLD mutant by immunoblotting. Consistent with drastic decrease in Rad53 activation, the degradation of Sml1 protein was found to be substantially decreased in the LD mutants in MMS treatment conditions compared to the wild type cells (Figure 2B, S2D-F). This impaired degradation of Sml1 protein probably limits Rnr1 availability, thereby may decrease the dNTP pool affecting DNA replication and repair.

Simultaneously, we also accessed the transcriptional response of Rnr genes. The basal *RNR3* expression levels were found to be significantly elevated in nLD mutants than the wild type cells (Figure S2I). This basal upregulation of *RNR3* reflects a deregulated state which is consistent either with underlying genomic stress or with non-specific degradation of its transcriptional repressor. However, following the MMS treatment, both wild type and H1246 mutant cells exhibited upregulation of *RNR2* and *RNR3*, however the magnitude of this induction was significantly much less in H1246 mutant than the wild type cells which correlates with the defect in Rad53 activation and Sml1 degradation (Figure 2C, S3A-B). The reduced *RNR* levels in the H1246 mutant may lead to decreased dNTP pool and inefficient DNA repair.

To further dissect the checkpoint defect in the LD deficient mutants, we next examined the activation status of Mec1, the upstream activation kinase responsible for initiating the activation of Rad53 kinase cascade as well as histone H2A phosphorylation. Consistent with the MMS driven Rad53 phosphorylation observed in wild type cells, total Mec1 levels increased progressively upon MMS treatments accompanied by a corresponding increase in γ-H2A phosphorylation (Figure 2D, S2J-M). In addition to monitoring the total Mec1, we also assessed the phosphorylation of Mec1 at S1991 site known to be phosphorylated by activated Rad53 which is indicative of the feedback regulation between Mec1 and Rad53 (22,23). The wild type cells showed increase in Mec1 S1991 phosphorylation levels, signifying efficient forward (Mec1-Rad53) and feedback (Rad53-Mec1) signalling of the checkpoint pathway. In contrast, LD deficient mutants (H1246 and Q47) exhibited reduced total Mec1 and severely diminished Mec1-S1991 phosphorylation (Figure S2J, S2L). Decrease in total Mec1 levels and S1991 phosphorylation in LD deficient mutants fails to robustly activate Mec1 signalling leading to significant decrease in Rad53 phosphorylation. In line with this, γ-H2A phosphorylation, which relies on Mec1 activity was found to be substantially decreased in LD deficient mutant, which further indicates that mutant exhibit defect in upstream DNA damage sensing mechanism (Figure 2E, S2G-H).

Taken together these finding demonstrate that LD deficient mutants fail to mount a proper checkpoint response upon MMS exposure. Impaired Mec1 activation, decrease in Rad53 phosphorylation, altered Sml1 degradation and poor expression of Rnr genes in LD deficient mutants collectively indicate defect in DNA damage sensing and repair mechanisms. This defect provides a mechanistic basis for the resistant phenotype of the LD deficient mutants demonstrating that their response arises from impaired sensing or subsequent relay of direct DNA damage signals.

### Neutral lipid droplet deficient mutants show elevated Doa1 expression and higher ubiquitination

Given that genes associated with neutral lipid droplet synthesis are not directly connected to canonical DDR pathway, we next investigated how the loss of lipid droplet inhibit the activation of checkpoint signalling. To address this, we examined known genetic factors whose deletion confers sensitivity to MMS and UV treatments. A previous study by Hanway et.al. reported six such genes, Doa1, Esc4, Tim13, Lge1, Rad33 and Rtt107 whose loss increases sensitivity to MMS and UV treatment conditions (24). Among these, only Doa1 deletion causes higher sensitivity to both MMS and UV. Doa1 is a ubiquitin chain binding regulatory protein of Cdc48 proteasomal regulatory axis (25,26). It has been also reported that Doa1 deletion leads to enhanced Rad53 phosphorylation (24). Since we previously observed resistance under MMS and UV and reduced Rad53 phosphorylation in LD deficient mutants under MMS treatment condition, we hypothesised that the LD mutant likely has elevated expression of Doa1. To test this possibility, we quantified the *DOA1* expression in wild type and the H1246 lipid droplets deficient mutant cells under increasing MMS treatments. RT-qPCR analysis revealed significantly higher expression of *DOA1* in the H1246 mutant in untreated and upon MMS treatment (Figure 3A, S3A-B). Studies have shown that the Doa1 overexpression disrupts its substrate selectivity and promotes non-specific ubiquitin chain linkage leading to proteostasis stress (27). To examine whether the elevated Doa1 expression in the H1246 mutants influence the ubiquitin profile, we examined global ubiquitination levels through immunoblotting using anti-ubiquitin antibody (Figure 3B). The H1246 mutant displayed marked increase in ubiquitinated proteins and corresponding reduction in free ubiquitin pool across all treatment conditions relative to wild type cells.

**Figure 3:**
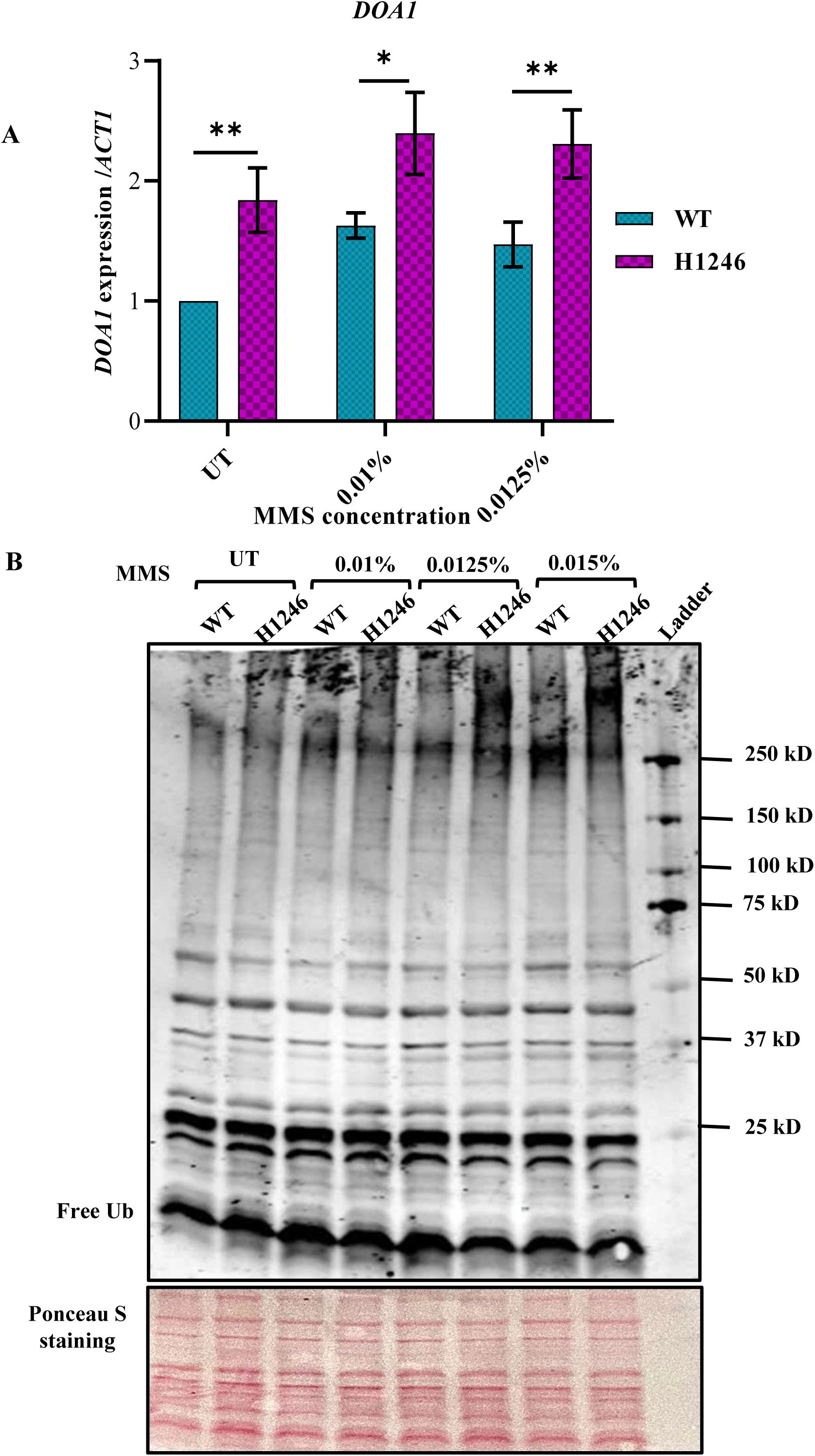
Increased *DOA1* expression and elevated ubiquitination levels in H1246 cells. **(A)** Expression level of *DOA1* was measured relative to *ACT1* mRNA in wild type and H1246 lipid droplet deficient mutant cells using RT-qPCR. Cells were grown in SC media till the late log phase and equal O.D. of cells were harvested to extract RNA using the hot phenol method. The cDNAs were synthesized using 400 ng of RNA, from the total RNAs of each strain. The data represents the mean of three independent biological replicates with the error bars depicting SEM (standard error mean). Statistical significance was calculated using Student’s t-test. **(B)** Western blot showing global ubiquitination levels of wild type and H1246 mutant in untreated and upon MMS treatment conditions. Ponceau-S staining of the probed blot is shown as protein loading control. Experiments were performed a minimum of three times (three independent biological replicates), and the representative image is shown here.

These results suggests that the elevated *DOA1* levels in the H1246 mutant correlate with enhanced and potentially dysregulated protein ubiquitination homeostasis. We believe that ubiquitination imbalance in the LD mutant may destabilize key regulatory kinases including those involved in DDR signalling which may be responsible for defects in sensing DNA damage and checkpoint activation. The constitutively higher *DOA1* expression and ubiquitination observed in the H1246 mutant may also be responsible for slower growth phenotype of this mutant. The excessive non-specific ubiquitination in the droplets deficient mutant probably increases the damaged protein pool suppressing the overall cellular fitness and the growth.

To further probe whether ubiquitination overload contributes to the MMS resistant phenotype of the H1246 mutant, we employed EGCG (Epigallocatechin gallate), a reported E3 ligase inhibitor (28).

We assessed the cell growth under MMS alone and MMS, EGCG co-treatment conditions. While the EGCG co-treatment did not substantially alter MMS sensitivity in wild type cells, the H1246 mutant displayed modest reduction in growth under MMS and EGCG co-treatment (Figure S3F, 3G). Strikingly, treatment with EGCG alone significantly improved the growth of H1246 mutant in liquid culture, whereas wild-type cells remained unaffected (Figure S3H). Together these findings suggest that the nLD deficient mutants exhibit chronic proteotoxic stress due to non-specific ubiquitination of proteins. We propose that the slow growth phenotype of the nLD deficient mutants is due to the drastic decrease in global protein pool which suppresses the overall fitness.

### Lipid droplets absence promote imbalance in sphingolipid/ceramide exchange

Next, we conducted a series of experiments to understand the link between Doa1 overexpression and lipid droplet absence. Previous studies have shown that perturbation of de-novo sphingolipid synthesis can modulate *DOA1* expression. Another study further demonstrates that exogenous supplementation of phytosphingosine (PHS), leads to hyperactivity of Doa1 which was determined by assessing the degradation of reporter substrate, Deg1-βGal (27). With this premise, we hypothesised that although the H1246 mutant has no direct perturbation in the sphingolipid biosynthesis, the absence of lipid droplets might rewire sphingolipid metabolism in a manner that functionally impacts their cytosolic presence. The altered sphingolipid pool may affect the Doa1 activity in the lipid droplet deficient mutant. To test this hypothesis, we first examined the expression of genes involved in the regulation of dynamic exchange between ceramide and sphingolipid bases as these two pools play essential role to maintain the sphingolipid homeostasis. Sphingolipid biosynthesis is initiated by condensation of palmitoyl-CoA with serine, a reaction catalysed by *LCB1, LCB2* and *TSC3*, resulting in formation of 3-ketohydrosphingosine. This intermediate is subsequently reduced by TSC10 to produce dihydrosphingosine (DHS) (29). The route of DHS formation is highly conserved in the higher eukaryotes and *Saccharomyces cerevisiae.* However, DHS onwards, yeast specific divergence defines distinct metabolic fates (Figure 4A) (30).

**Figure 4:**
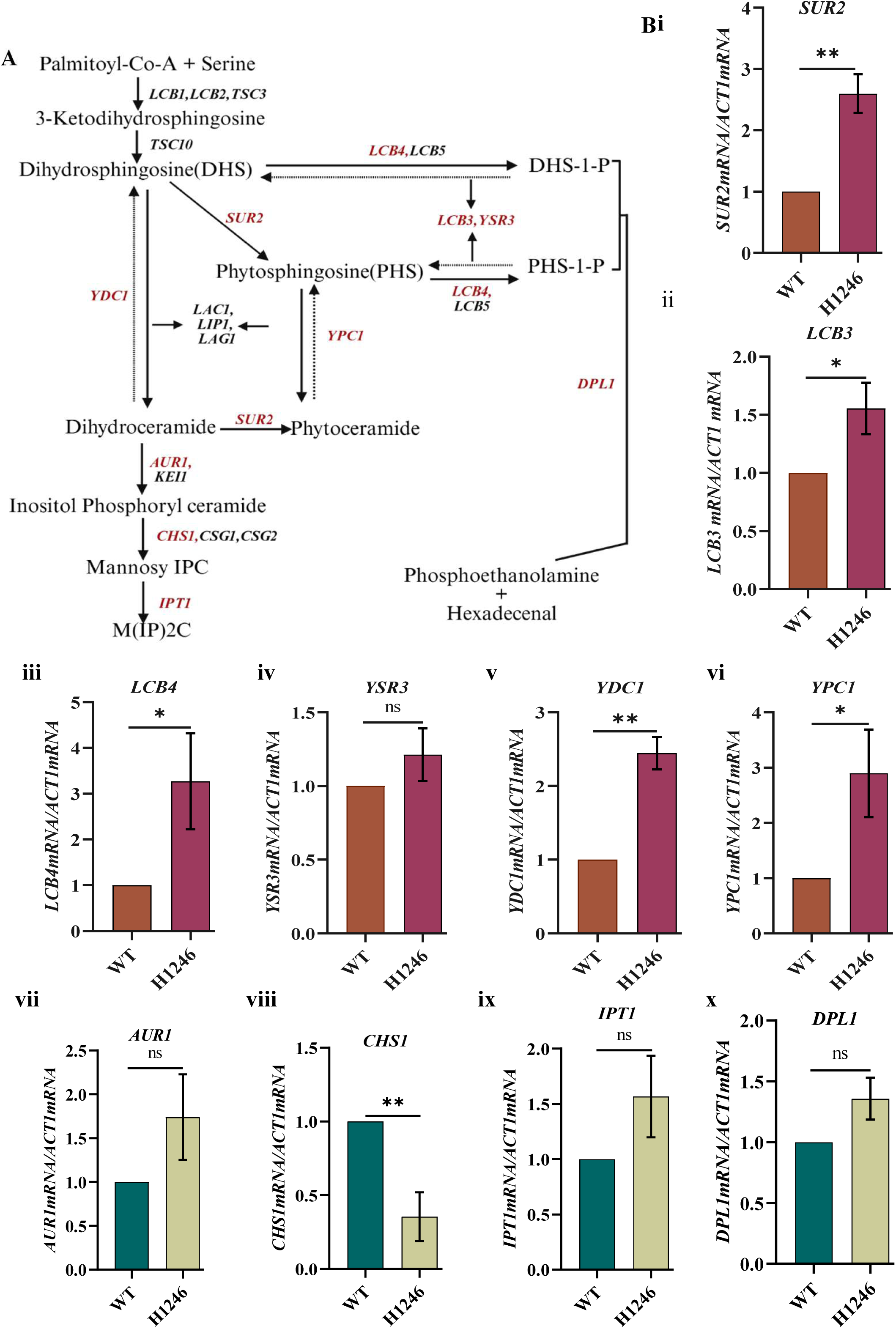
Lipid droplet deficiency enhances the exchangeable sphingolipid pool in H1246 mutant. **(A)** Flowchart to show the sphingolipid biosynthesis pathway beginning from Palmitoyl-Co-A and serine condensation. **(Bi-Bvi)** RT-qPCR to measure the m-RNA expression levels of different enzymes that regulate the dynamic exchange between ceramide and complex sphingolipids. Expression levels were measured relative to *ACT1* mRNA in wild type and H1246 mutant strains using RT-qPCR. Cells were grown in SC media till the late log phase and equal no. of cells were harvested to extract RNA using the hot phenol method. From the total RNAs of each strain, 400ng was used to synthesize cDNA. **(Bvii-Bx)** RT-qPCR to measure the expression levels of *AUR1, CHS2, IPT1* and *DPL1* genes encoding the enzymes that channel ceramide/ sphingolipid intermediates out of the exchangeable pool by converting them into terminal or non-recyclable products. The data represents the mean of three independent biological replicates with the error bars depicting SEM (standard error mean). Statistical significance was calculated using Student’s t-test.

In yeast cells, the DHS can either be converted into DHS-1-P by *LCB4* and *LCB5* or to dihydroceramide regulated by *LAC1* and *LAG1* which encodes ceramide synthases. All these reactions are reversible, and maintains a dynamic balance between sphingolipid bases and ceramide (31,32). To understand, how neutral lipid droplets deficiency influences this complex network, we quantified the gene expression of the key conversion enzymes. To this end, we found higher expression of *SUR2, LCB3, LCB4, YPC1,* and *YDC1* in the H1246 mutant strain which may lead to increase in phytosphingosine and dihydrosphingosine concentration in the cells (Figure 4Bi-vi). We further checked the expression of genes involved in non-reversible ceramide drainage pathways such as *AUR1, CHS2, IPT1* and *DPL1* which regulates the non-reversible flux to decrease the ceramide from the exchangeable pool. *AUR1*, *CHS2* and *IPT1* regulates the conversion of dihydroceramide to complex mannosyl-(Inositol-phosphoryl)_2_ ceramide whereas *DPL1* regulates the conversion of DHS-1-P and PHS-1-P to phosphoethanolamine and hexadecenal. We did not detect any significant change in the expression of *AUR1*, *IPT1* and *DPL1* in the H1246 mutant, but the expression of *CHS2* was found to be markedly reduced suggesting a reduced capacity of the mutant to funnel ceramide away from exchangeable pool (Figure 4Bvii-x).

Sphingolipids and ceramides exist in a reversibly exchangeable pool, where each can be interconverted depending on metabolic demand. When ceramides accumulate in excess, they are neutralized through O-acylation by two LD biosynthesis genes, dga1 and lro1. Thereby the excess ceramides gets diverted into neutral lipid droplets(Figure 5A) (33). To test whether elevated sphingolipid bases themselves can influence LD abundance, we supplemented cells with exogenous PHS and monitored LD number under these conditions. The wild type cells were treated with 20µM PHS followed by staining with BODIPY (493/503nm) to visualize lipid droplets. Given that lipid droplet synthesis enhances under stress conditions, we also examined the effect of PHS supplementation on the growth of wild type and the H1246 mutant strain by spot test assay. We observed that supplementation of PHS does not affect the growth of cells (Figure 5C, S4C). Interestingly, the microscopic analysis revealed that PHS supplementation increases the number of lipid droplets in wild type cells, but we did not detect any BODIPY fluorescence in H1246 mutant (Figure 5B, S4A-B). For comparison, cells were also treated with MMS and MMS plus PHS which revealed a distinct pattern of lipid droplet distribution between these treatments. Together, these results indicate that the inability of lipid droplet deficient mutants to store sphingolipid and ceramides increases the cytosolic sphingolipid-ceramide pool. This lipid imbalance is probably responsible for the upregulation of Doa1 expression which triggers the excessive ubiquitination and proteostasis dysregulation in the LD deficient cells (Figure 5D).

**Figure 5:**
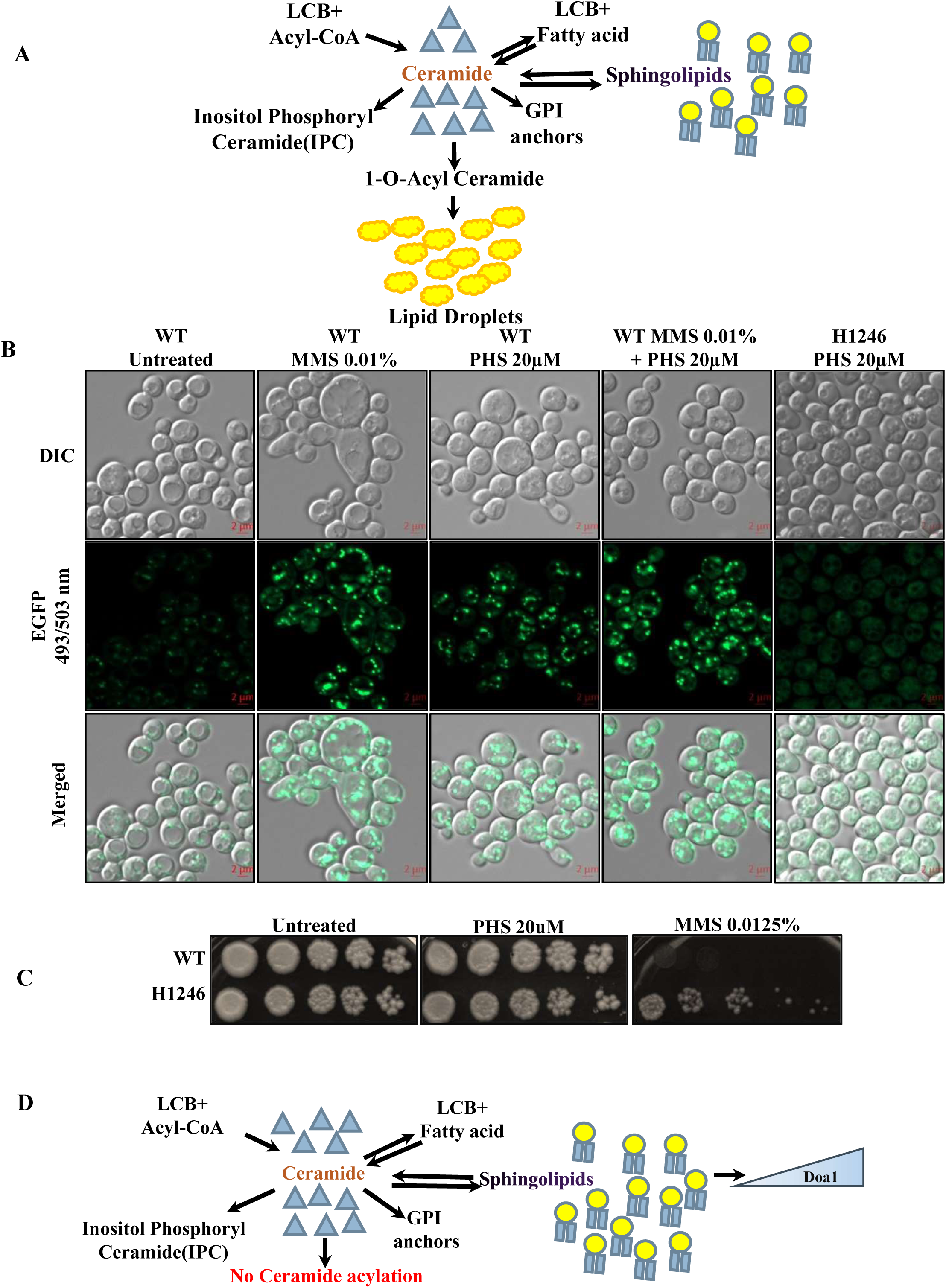
Excess cytosolic PHS enhances lipid droplet synthesis. **(A)** Schematic representation of ceramide and sphingolipid reversible conversion and their storage in lipid droplets. **(B)** Microscopy using BODIPY stain indicates increase in the lipid droplet synthesis upon external PHS supplementation in wild type cells. Cells grown till exponential stage were treated with PHS 20µM and 0.01% MMS for 2hours and then stained with BODIPY (493/503) and images were captured using Zeiss-Apotome microscope. **(C)** Spot test assay to examine the effect of PHS treatment on the growth of wild type and H1246 mutant strain. Equal volumes of each culture from 5-fold serial dilutions of initial O.D._600_∼1 were spotted in SC agar plates containing 20µM PHS and 0.0.1% MMS and incubated for 2-4 days. Images were captured by HP scanner. SC plate without any treatment was spotted as control plate. Experiments were performed a minimum of three times (three independent biological replicates), and the representative image is shown here. **(D)** Schematic representation of alterations in ceramide and sphingolipid pool arise due to absence of lipid droplets. Increased cytosolic sphingolipids might trigger the Doa1 dependent protein ubiquitination and no-specific protein degradation.

### Persistent DDR defect drives genome instability and premature ageing

We next examined, how does compromised DNA damage sensing affect long-term genome integrity and cellular ageing. Consistent with previous reports the chronological lifespan of lipid droplet deficient mutant is very small (3-4 days). We also observed normal 9 days long chronological lifespan of the wild type cells, however, the quadruple lipid droplets deficient mutants failed to grow beyond the third day, indicating a markedly decrease in the lifespan (Figure S5A-B).

Primary ageing hallmarks are compromised DNA (nuclear and mitochondrial) integrity and accumulation of mutations. To check DNA integrity, we monitored γ-H2A levels (phosphorylation at serine 129) as readout of the accumulated DNA damage (34,35). In the wild type cells, γ-H2A levels remained largely constant over 12-15 days whereas H1246 mutant showed a steep and sustained increase in γ-H2A phosphorylation from the third day onwards, which displays the higher accumulation of DNA damage (Figure 6A-B). The total proteins separated through the SDS-PAGE gel was stained with Coomassie dye to ensure that apparent increase in γ-H2A phosphorylation in the lipid droplet deficient mutant is not due to unequal loading. Despite equal loading of proteins, confirmed by Coomassie staining, the protein samples of H1246 mutant showed altered α-Tbp signal intensity analysed by western blotting. The Coomassie blue dye-stained protein gel revealed almost equal intensities of protein bands between wild type and the mutant towards high-molecular weight range, but a visible smear and variable levels of protein bands were observed towards lower molecular weight range in the samples of H1246 mutant, indicative of poor quality of proteins which may be due to proteostasis stress and not due to the loading errors.

**Figure 6:**
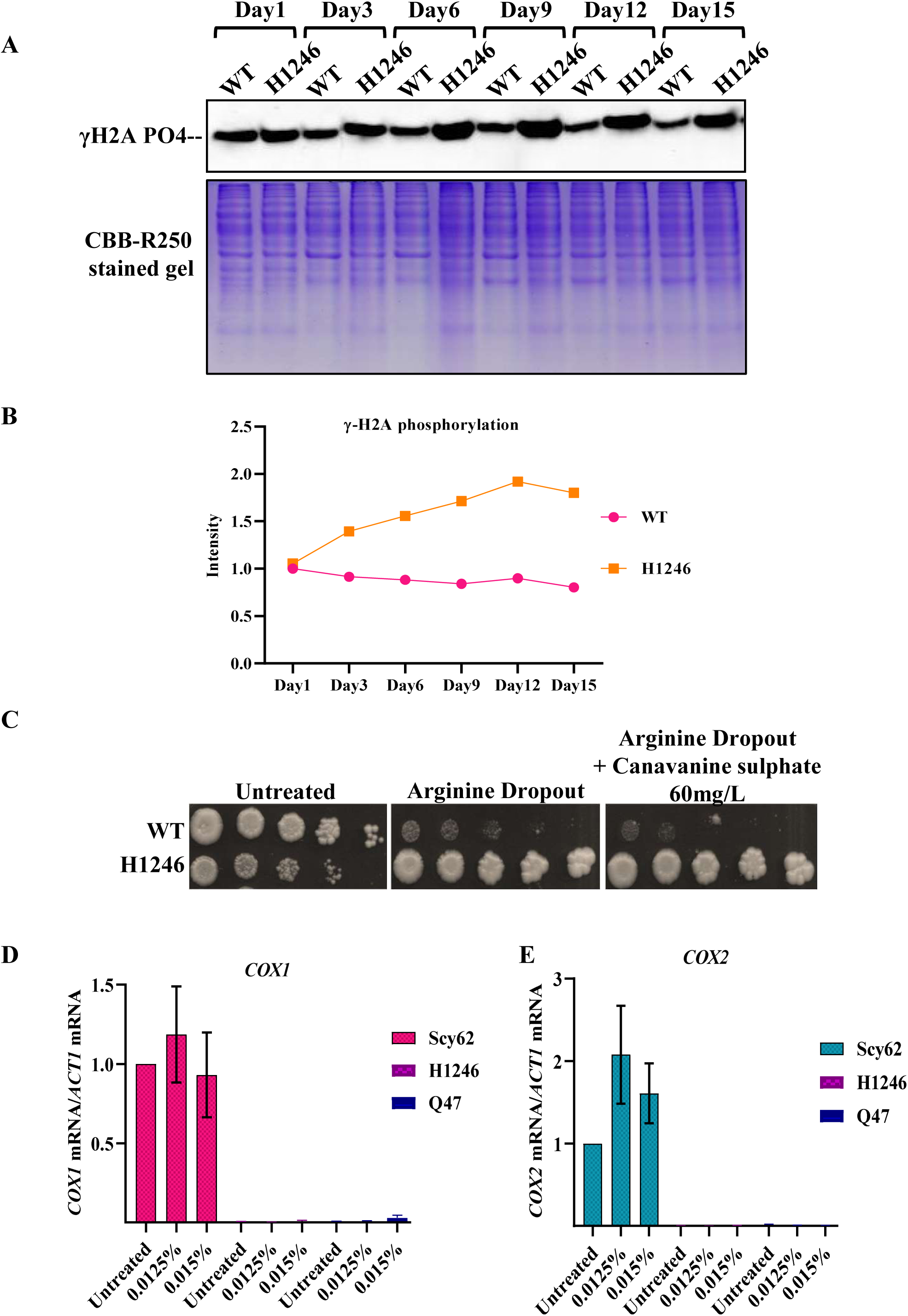
Persistent defect in DNA double strand break repair result into significant increase in γ-H2A phosphorylation, higher mutation burden and decreased chronological lifespan. **(A)** Spot assay of wild type and H1246 mutant cells at different days from continuously growing cultures to monitor the chronological life span. Wild type cells show healthy growth while the H1246 mutant shows complete loss of viability after day 3. **(Bi)** Western blot to compare the γ-H2A accumulation between wild type and H1246 mutant cells at different days of chronological life span. Cells were continuously grown at 30°C/200rpm for 15 days and 10 O.D. of cells were taken at depicted time point to prepare whole cell extract. Coomassie blue stained gel and α-TBP western blot performed for the loading control. **(Bii)** Quantification of presented blot shown relative to α-TBP probing. **(C)** Spot test showing effect of canavanine resistance of H1246 strain due to mutation in arginine permease activity. Equal volumes of cells from 5-fold serial dilutions of initial O.D._600_∼1 was spotted in synthetic arginine dropout growth media with and without 60mg/l canavanine sulphate and incubated for 2-4 days. Images were captured by HP scanner. Cells were also spotted on synthetic complete agar media for the control. Experiment was performed a minimum of two times (two independent biological replicates), and the representative image is shown here. **(D)** Expression level of *COX1* and *COX2* was measured relative to *ACT1* mRNA in WT, H1246 and Q47 strains using RT-qPCR. The data represents the mean of three independent biological replicates with the error bars depicting SEM (standard error mean). Statistical significance was calculated using Student’s t-test.

Persistent DNA damage not only promotes mutation in the nuclear genome but also jeopardise mitochondrial DNA integrity. Mitochondrial DNA, being unprotected by histone proteins, is particularly more vulnerable to oxidative and genotoxic stress exposures (36). We therefore quantified the expression of mitochondrial genes *COX1* and *COX2*, which we found nearly absent in lipid droplet deficient mutants (H1246 and Q47) even under untreated condition (Figure 6D-E). In further experiments, we found that H1246 mutant is unable to grow on SCEG (Ethanol and glycerol, non-fermentable Carbon source)(Figure 7A). Together these finding suggests that due to compromised DDR response the mitochondrial DNA as well as function is damaged in the LD deficient mutants.

**Figure 7:**
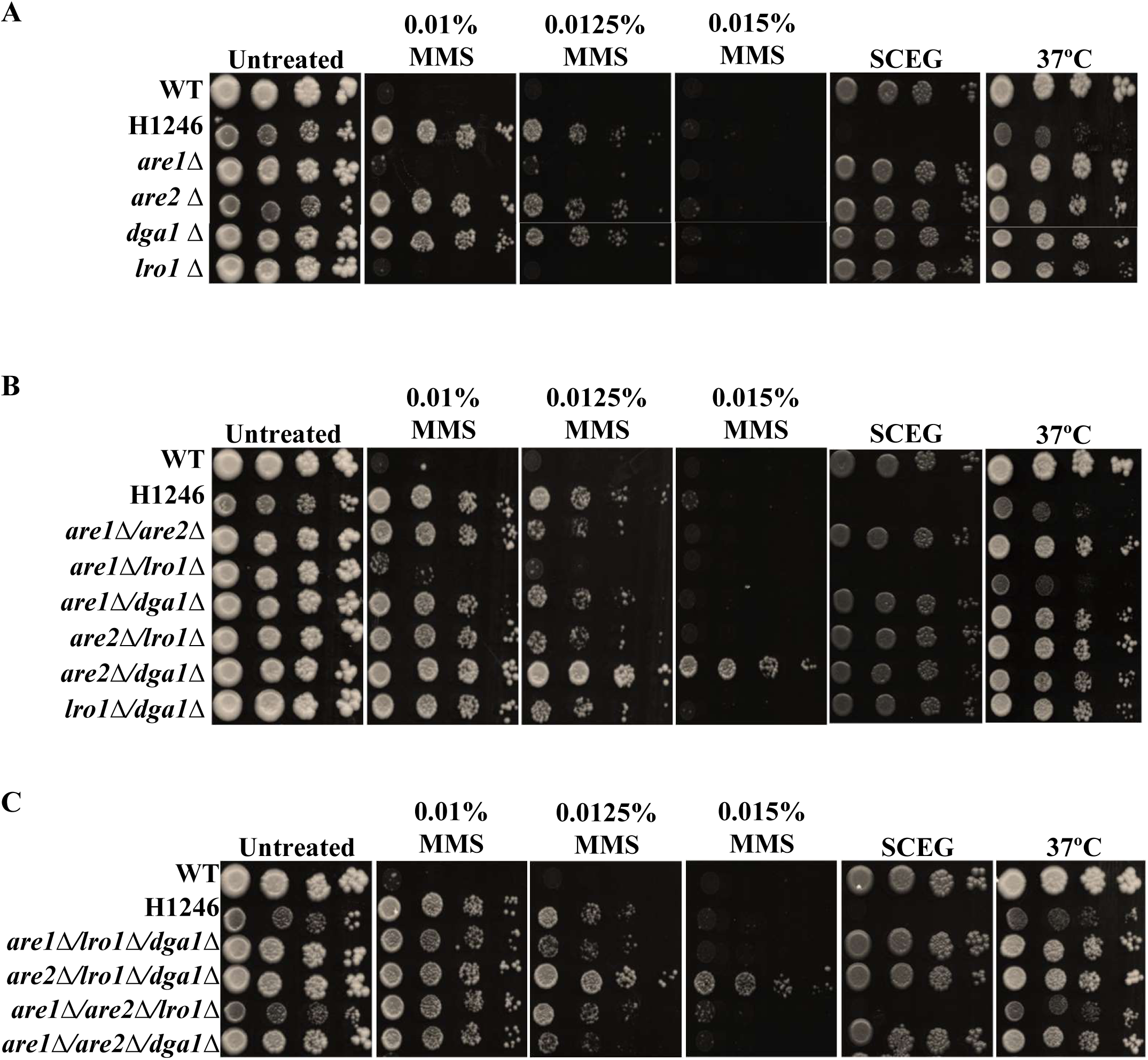
Combinatorial gene deletion analysis reveals the role of lipid droplets in DNA damage response and mitochondrial functions. **(A)** Spot assays to examine the growth of are1Δ, are2Δ, lro1Δ and dga1Δ in MMS, SCEG media and at 37°C. Deletion of genes was created using a standard method based on homologous recombination. Growth on SCEG media was used to monitor mitochondrial health needed to metabolize the non-fermentable carbon sources. **(B, C)** Spot test assay to monitor the growth of combinatorial double and triple deletion mutants of are1Δ, are2Δ, lro1Δ and dga1Δ on MMS, SCEG media and at 37°C.

Further we assessed the effect of compromised RNR expression and damage repair on mutation accumulation. We performed a canavanine resistance assay, a classical test for assessing mutation frequency in yeast cells (37). The assay depends upon the activity of the *CAN1* gene which encodes an arginine permease that also transports toxic arginine analogue canavanine. The wild type cells (functional *CAN1*^+^) import canavanine and die whereas mutants with loss-of-function mutations in CAN1 (*can1*^-^) survive and become resistant to canavanine (Can^R^). When we spotted wild-type and nLD cells on arginine deficient media which contains canavanine, the H1246 mutant displayed robust growth on canavanine compared to wild type cells, indicating higher spontaneous mutation rates in the mutant due to chronic DDR failure (Figure 6C, S5C). The growth of wild type cells decreases in arginine dropout plate compared to SC plate and addition of canavanine show severe toxic effects.

Taken together, these findings demonstrate that the loss of lipid droplets leads to failure of sensing and repair of the DNA damage causing accumulation of mutations. Higher frequency of mutations in the LD deficient mutant results in nuclear and mitochondrial genomic instability which accelerate the ageing process. Compromised mitochondrial function in LD deficient mutants in not only due to alteration in sterol metabolism and channelling (as reported previously) but also due to mitochondrial DNA damage which eventually compromises the function.

### Genetic screening of neutral lipid droplet synthesis shows differential contribution of individual genes to DNA damage response

To understand whether the growth phenotypic defects observed in the H1246 mutant arise from combined loss of all four lipid droplet biosynthesis genes (are1Δare2Δlro1Δdga1Δ) or a particular gene is responsible, we generated single and combinatorial double and triple deletion mutants of these lipid droplet biosynthesis gene (38). The growth of these mutants was assessed under MMS treatment, elevated temperature (37°C) and on non-fermentable carbon source containing growth media (SCEG) to study their DDR response, proteostasis and mitochondrial function, respectively (39,40). Among the single deletion mutants, are2Δ and dga1Δ mutants exhibit resistant growth phenotype same as H1246 mutant, whereas are1Δ and lro1Δ single deletions were highly sensitive, showing poor growth even at a very low concentration (0.01%) of MMS (Figure 7A, S6A). Growth curve analysis in SC liquid growth media clearly shows that are1 and lro1 deletion mutants exhibit similar growth pattern like wild type cells whereas the are2 and dga1 deletion mutants show similar tolerance phenotype towards MMS treatment as H1246 mutant (Figure S6F, H). None of the single gene deletion mutant show defect on SCEG media, suggesting that single deletions do not affect mitochondrial health. Similarly, their growth at 37°C was found to be same as wild type cells.

The double deletion mutant, are2Δ/dga1Δ displayed enhanced resistance compared to the H1246 mutant as well as are2Δ and dga1Δ strain, suggesting a synergistic effect between these two genes. Interestingly, are1Δ/lro1Δ mutant showed slightly better growth than the wild type cells, despite the at par sensitivity of their single deletion mutants to MMS (Figure 7B, S6B). All the triple deletion mutants showed resistant to MMS same as H1246 mutant. However, among all these triple deletion mutants the are2/lro1/dga1 mutant where only Are1 is present, showed much higher resistance to MMS.

The combinatorial deletion also highlighted the regulatory cross talk between the LD biosynthesis genes to maintain mitochondrial health. We observed that although deletion of one gene is not sufficient to affect mitochondrial function, are1 and lro1 has specific interaction to support mitochondrial function. The double deletion mutant are1Δ/lro1Δ and triple deletion mutant, are1Δ/are2Δ/lro1Δ showed no growth on SCEG plate similar as H1246 strain (Figure 7B and 7C, S6B and S6C). double and triple deletion mutants with defective mitochondrial health also showed sensitivity towards elevated temperature (37°C). The mitochondrial defect persist when Are2 is deleted, so the loss of are2 neither supresses nor prevents the defect. In contrast, deleting dga1 together with are1 and lro1 supresses mitochondrial defect. Together these results underscore a complex functional network among the genes that regulate the metabolism of neutral lipids playing essential roles in DNA damage repair signalling pathways and mitochondrial health.

To further understand how lipid droplet dynamics and Rad53 activation are influenced under MMS induced DNA damage stress, we selected two contrasting mutants based on their MMS sensitivity phenotype for further analysis, the are1Δ mutant most sensitive and are2Δ/lro1Δ/dga1Δ mutant most resistant (Figure S6D, S6E). We, first the examined the neutral lipid droplets accumulation using BODIPY (493/503nm) staining (Figure 8A, S7A-B) (4). In untreated condition, are1Δ cells exhibit similar level of fluorescence as the wild type, however, upon MMS exposure, fluorescence intensity markedly increases, indicating elevated neutral lipid droplet synthesis. Interestingly, in are1Δ cells, the lipid droplets appeared smaller and numerous compared to wild type cells, where larger, patch like accumulation were observed. This observation implies that are1 deletion alters the size, remodelling and distribution of lipid droplets under stress conditions.

**Figure 8:**
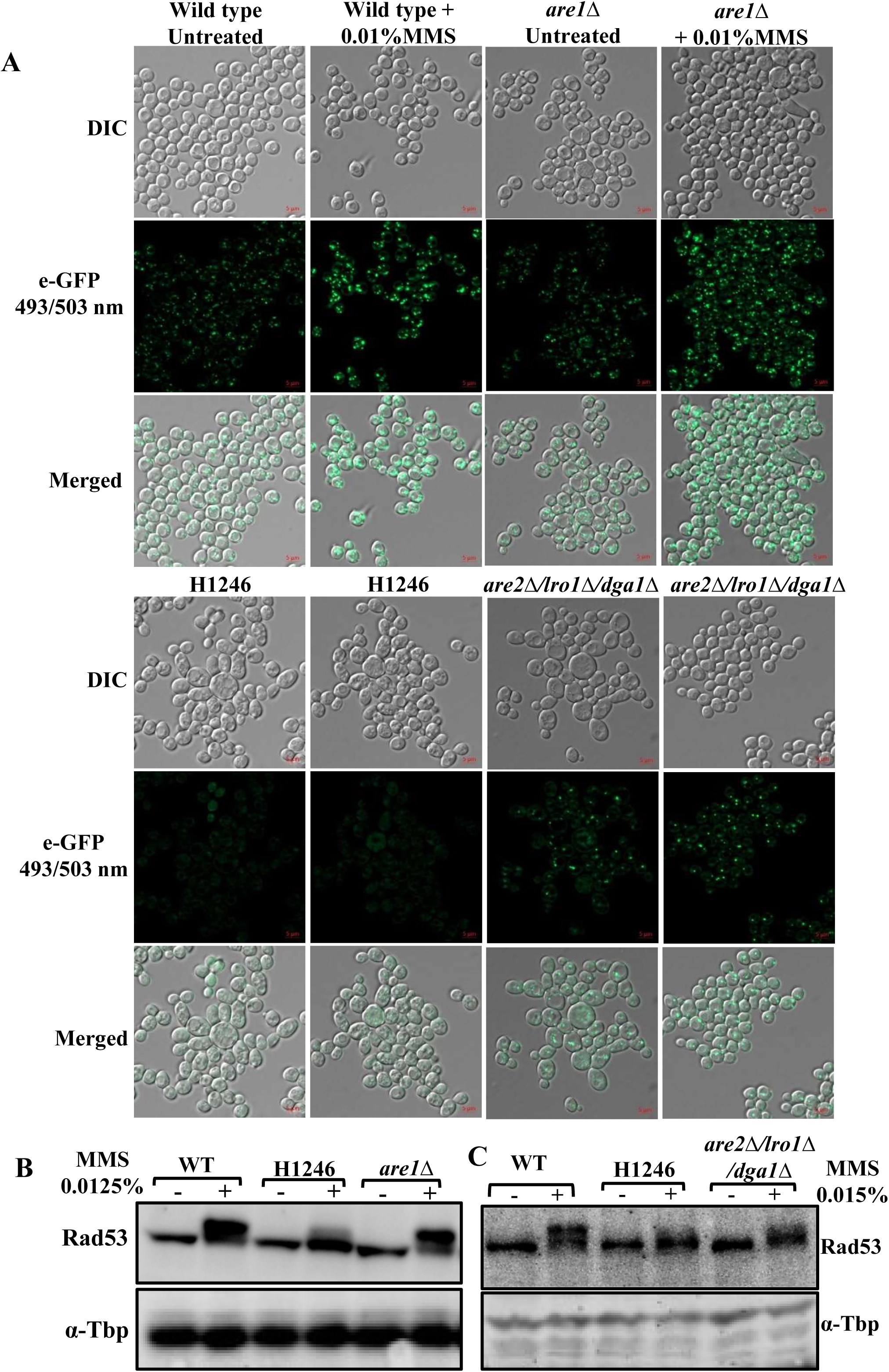
Are1 mediated regulation on Rad53 phosphorylation and lipid droplet biogenesis. **(A)** Microscopic analysis of lipid droplets distribution by staining with BODIPY (493/503) in wild type, H1246 lipid droplets deficient mutant, are1Δ and are2Δ/lro1Δ/dga1Δ mutants in untreated and 0.01% MMS treatment. Triple deletion mutant, are2Δ/lro1Δ/dga1Δ show significantly low lipid droplet intensity in untreated and upon MMS treatment. **(B)** Western blotting to measure Rad53 phosphorylation in wild type, H1246 mutant and are1Δ in untreated and upon 0.0125% MMS treatment. In MMS treatment condition, are1Δ strain showed very high Rad53 phosphorylation. Western blotting with α-Tbp antibody was performed for protein loading control. **(C)** Western blotting to measure Rad53 phosphorylation in wild type, H1246, and in are2Δ/lro1Δ/dga1Δ mutant upon 0.015% MMS treatment for two hours. At 0.015% of MMS treatment, H1246 and are2Δ/lro1Δ/dga1Δ mutants strain showed drastic decrease in Rad53 phosphorylation compared to wild type cells. Western blotting with α-Tbp antibody was performed for protein loading control. Western blot and microscopy experiments were performed three times (three independent biological repeats), only the representative images are presented here.

In contrast, are2Δ/lro1Δ/dga1Δ triple mutant showed a striking reduction in overall lipid droplet content, with only sparse and enlarged droplet detectable across the population. Upon MMS treatment, this triple deletion mutant strain exhibited only a slight increase in lipid droplet number, suggesting that lipid droplet biogenesis is impaired in this mutant. Further, we analysed the Rad53 phosphorylation under MMS treatment to probe the alterations in the DNA damage response due to change in lipid droplet biosynthesis. Interestingly, we observed increase in Rad53 phosphorylation upon MMS treatment in are1Δ mutant cells. The higher Rad53 phosphorylation in are1Δ mutant cells in comparison to the wild type cells correlates with their higher sensitivity to DNA damage (Figure 8B, S7C-E). Conversely, in comparison to wild type cells, the are2Δ/lro1Δ/dga1Δ mutant showed significantly less phosphorylation of Rad53 same as quadruple mutant which correlates with their resistant phenotype to MMS treatment. Above results suggest that the absence of are1 (are1Δ) enhances and the presence of are1 (are2Δ/lro1Δ/dga1Δ) supresses the Rad53 phosphorylation under genotoxic stress (Figure 8C, S7F-G). These finding indicate that the lanosterol esterification enzyme are1 appears to modulate the DNA damage response signalling pathways through lipid droplet linked mechanism, and together with lro1 has supportive role for mitochondrial health which needs to be explored through detailed investigation of altered lipidome for deeper insight.

## Discussion

Lipid droplets (LDs) are ubiquitous, monolayer-bound dynamic organelles present across all eukaryotic cells. Structurally, they are consist of a hydrophobic core composed primarily of sterol esters and triacylglycerols, enclosed by phospholipid monolayer derived from the endoplasmic reticulum (ER) membrane (41). This unique monolayer distinguishes LDs from other double-membrane-bound organelles and enables a fluid interface for interactions with cytosolic components and proteins involved in lipid metabolism, trafficking, and cellular signalling(42). Beyond their canonical role as neutral lipid storage depots, LDs actively contribute to membrane synthesis, protein quality control, and energy homeostasis. Recent studies further reveal that LDs dynamically respond to diverse environmental and metabolic cues(12,43). Notably, their number and size increase markedly under oxidative and genotoxic stress, suggesting a protective and adaptive role. For instance, exposure to reactive oxygen species or DNA-damaging agents induces robust LD biogenesis in yeast, mammalian, and plant cells (43). However, despite these consistent observations, the precise functional contribution of LDs during genotoxic stress remains poorly understood. Addressing this gap is critical to uncovering how lipid metabolism interfaces with DNA damage signalling and repair pathways.

To investigate this, we examined how the absence of LDs influences cellular response to DNA damage. Growth analysis via spot assay and growth curve revealed an unexpected phenotype in LD-deficient mutants. Although previous reports suggest that LDs confer protection under genotoxic stress, implying growth arrest or sensitivity when absent, we observed improved growth of LD-deficient mutants following methyl methane sulfonate (MMS) and ultraviolet (UV) treatments (Figure 1A-B). This observation prompted us to examine checkpoint activation in wild type and LD-deficient mutants under untreated and DNA-damaging conditions. Immunoblot analysis showed that activation of the checkpoint sensor kinase Mec1 and its downstream effector Rad53 was significantly reduced in LD-deficient mutants. Correspondingly, phosphorylation of Mec1 and Mec1-dependent γ-H2A was constitutively low in these strains.

To understand the MMS- and UV-resistant phenotype of LD-deficient mutants, we searched for genes which upon deletion renders the cells sensitive to MMS and UV. A previous report showed *DOA1* deletion mutant to be hypersensitive to MMS and UV and exhibits increased Rad53 phosphorylation. We therefore hypothesized that *DOA1* may be overexpressed in LD-deficient mutants, thereby Rad53 phosphorylation is reduced. *DOA1* is functionally linked to the Cdc48-dependent proteasomal degradation pathway, influencing ubiquitin chain specificity and substrate turnover when dysregulated (44). In LD-deficient mutants, we observed elevated *DOA1* expression at basal as well as under MMS treatment conditions. Studies indicate that components of the Mec1-dependent checkpoint are subject to ubiquitin-mediated regulation, where K48-linked ubiquitin chains promote proteasomal turnover and K63-linked chains facilitate repair factor recruitment (26). Higher Doa1 expression correlates with enhanced ubiquitin-mediated protein degradation, destabilizing protein which are crucial for cellular fitness and stress adaptation which we also observed in H1246 mutant upon probing global protein ubiquitination, In addition to the global imbalance observed in H1246 cells, we observed an unexpected feature related to Atg8 associated processing. During the Atg8-GFP immunoblot analysis, we consistently detected a ∼75kDa reactive band in wild type cells which likely represents an Atg8 conjugated substrate, given that Atg8 is fused to GFP. Notably this band remains unaffected upon rapamycin and cerulenin treatment in wild type cells indicating that although that substrate is Atg8 modified, it does not undergo processing during canonical autophagic flux. Importantly, none of the LD biosynthesis genes are known Atg8 substrates. We observe that ∼75kDa band was completely absent in all H1246 samples, irrespective of treatment (Figure S3C-E). Given our overall observation that Doa1 is overexpressed in H1246 cells and Doa1 modulates Cdc48 dependent proteasomal extraction and degradation, it is plausible that this Atg8 modified species become destabilizes specially in mutants. In wild type cells this substrate appears refractory to autophagic processing and yet in H1246 cells, the hyperactivated Doa1-Cdc48 axis may target it aberrantly, leading to non-specific degradation (45). Although this observation emerged serendipitously and identity of the protein remains unknown, the complete loss of this otherwise stable Atg8-modified species provide additional supporting evidence for deregulated Cdc48 guided degradation in H1246 (46).

Altogether, we propose that Doa1 overexpression in LD-deficient cells shifts the proteasomal balance toward increased K48-linked degradation of crucial cell proteins which reduces their stability and impairs stress signalling (47). Although LDs have been implicated in ubiquitination regulation, the specific link to Cdc48-dependent degradation via Doa1 has not been explored.

The next question was to understand how Doa1 hyperactivity is related to absence of Lipid droplets? Doa1 activity is modulated by the levels of phytosphingosine (PHS) and under normal conditions, excess ceramide is neutralized through O-acylation by Dga1 and Lro1, thereby maintaining lipid homeostasis and thus the proteostasis (33) (Figure 5A). However, in LD-deficient mutants, this neutralization route is disrupted, leading to cytoplasmic accumulation of bioactive sphingolipids such as PHS. This imbalance likely hyperactivates Doa1, driving excessive ubiquitination and proteasomal degradation, thus impairing DNA damage sensing. Supporting this idea, we observed upregulation of genes including *SUR2*, *LCB3*, *YPC1*, and *YDC1*, all involved in diverting ceramide flux toward phytosphingosine in the H1246 LD-deficient strain. Additionally, increased LD count in wild-type cells upon external supplementation of PHS show that LDs act as the storage site for neutralized excess sphingolipid/ ceramides. Together, these findings suggest that LDs act as buffering reservoirs that maintain sphingolipid equilibrium, preventing hyperactivation of lipid-sensitive ubiquitin regulators such as Doa1 and thereby preserving genomic stress signalling fidelity (Figure 5D).

Moving forward, the consequences of impaired DNA damage signalling on ageing hallmarks were studied on LD deficient mutants. Persistent defects in DNA sensing and repair evidenced by elevated γ-H2A levels, indicates accumulation of unrepaired DNA lesions. This chronic DNA damage burden correlated with decreased survival in chronological aging assays. These observations suggest that defective DNA repair contributes to the shortened lifespan of LD-deficient mutants. Analysis of mitochondrial integrity revealed reduced expression of *COX1* and *COX2*, indicative of mitochondrial DNA damage, potentially accounting for compromised respiratory function (Figure 6D and 6E). Thus, loss of LDs disrupts maintenance of both nuclear and mitochondrial genomes, promoting premature aging (45). Previous reports have shown that disruption of neutral lipid storage alters sterol metabolism and impairs mitochondrial function (40). Our results further suggest that LD-deficient cells accumulate more mutations across generations due to defects in DNA damage sensing and repair, leading to genomic instability, damage of mitochondrial genome and function. Together, these health alterations contributes to accelerated ageing (Figure 9).

**Figure 9:**
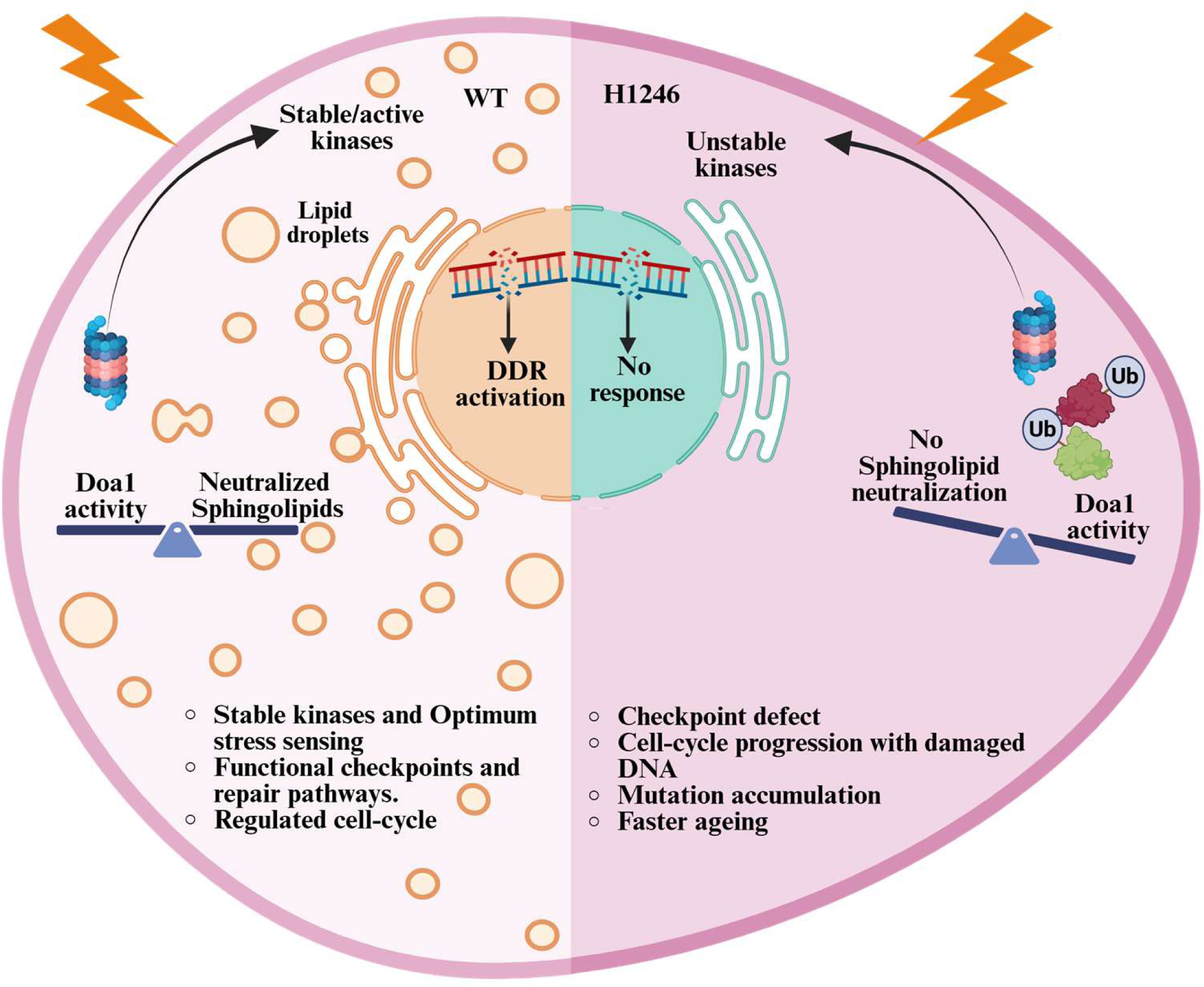
Proposed model illustrating on how the loss of neutral lipid droplets disrupts sphingolipid homeostasis, Doa1 activity, proteostasis, and DNA damage response. The biosynthesis of neutral lipid droplets balances sphingolipid–ceramide cycling in wild type cells. Adequate formation of lipid droplets maintains proteostasis, regulates the activity of sensory kinases that enables optimal stress sensing and activation of checkpoints, efficient DNA repair, and regulates cell-cycle progression. On the other hand, the absence of lipid droplets in the H1246/Q47 mutant cells results in the accumulation of free sphingolipids, which cannot be efficiently acylated and stored in lipid droplets. Elevated sphingolipid levels elevate Doa1 activity, causing disruption in proteostasis and increased proteasome mediated degradation of sensory kinases. Defect in upstream DNA damage sensing mechanism causes decrease in DNA double strand break repair response. As a result, lipid droplet deficient cells exhibit inefficient checkpoint activation, progress through the cell cycle with damaged DNA, accumulate mutations leading to accelerated cellular aging.

To further dissect genetic factors linking LDs to DNA damage signalling, we generated single and combinatorial deletion mutants of key neutral lipid biosynthesis genes (*are1*, *are2*, *dga1*, *lro1*) and examined their responses to DNA-damaging agents. Our data revealed a distinct role for *ARE1* in modulating DNA damage signalling: *are1*Δ strain exhibited enhanced Rad53 activation and increased DNA-damage sensitivity. In contrast, *are2*Δ/*lro1*Δ/*dga1*Δ mutant showed reduced Rad53 activation and increased resistance to DNA damage, like LD-deficient strains. Notably, *are1*Δ also exhibited an abnormal LD morphology, characterized by increased LD number but irregular cytoplasmic distribution following MMS exposure (48). These findings suggest a previously uncharacterized genetic interplay among neutral lipid synthesis genes in regulating stress responses. A comprehensive proteomic, lipidomic, and live-cell imaging analysis will be required to resolve these interactions.

Taken together, our study establishes a mechanistic link between lipid droplet biogenesis, sphingolipid homeostasis, and DNA damage sensing fidelity. We demonstrate that LDs safeguard cells from intrinsic lipid-mediated toxicity by regulating sphingolipid flux and proteostasis. Disruption of LD formation leads to an imbalance in sphingolipid metabolism and protein turnover, resulting in genomic instability and reduced chronological lifespan.

## Material and methods

### Media and strains

The yeast strains SCY62 (WT) and H1246, are kind gift from Professor Sten Stymne (4). Methyl methane sulphonates (MMS), hydroxyurea (HU), Epigallocatechin gallate (EGCG) and amino acid for synthetic media preparation are procured from Sigma-Aldrich. Reagents used in the present study are of molecular biology grade and purchased from Sigma-Aldrich, Himedia, KAPA biosystem, Invitrogen, Bio-Rad, and MP Biomedicals. SC growth media composed of 0.181 % SC dropout mix, 0.17 % yeast nitrogenous base without inositol (Himedia), 0.5 % ammonium sulphate (Sigma-Aldrich), and 2 % D-glucose (Sigma-Aldrich) (49). The yeast cells were grown at 30 °C, 200 rpm in the incubator shaker (Innova 44, New Brunswick Scientific). The SC agar plates were prepared by adding 2 % agar (Bacto agar, BD) powder into SC liquid growth media. The standard laboratory protocol was used to grow yeast cells (Yeast protocol handbook, Clon tech Laboratory, Inc). Cells were grown SC growth media in glass tubes or flasks at 30°C in incubator shaker at 200rpm. The cells were spotted on plates containing SC agar growth media were incubated for 3-4 days at 30°C in the incubator.

### Competent cell preparation and transformation

Homologous recombination method was used for gene deletion and primers were designed as per method described by Brachmann *et.al* using PRS plasmids (50). PCR products were synthesized using two-step PCR method and purified before transformation. Competent cells were prepared by lithium acetate method and after transformation, cells were spread on SC amino acid drop-out media. Selected colonies were restreaked on similar auxo-trophic selection plates and genomic DNA was extracted for PCR confirmation. List of strains used and generated in this study is given in supplementary table 3.

### Spot test or drop dilution test

To evaluate the growth fitness, we used growth sensitivity assay (spot assay) as described previously (51). Cells were grown overnight in synthetic complete media at 30°C/200rpm and were dilute 10-fold or 5-fold serially (10^-1^ to 10^-4^ or 5^-1^ to 5^-5^) and spotted on SC agar plate containing different chemical agents like MMS, HU, CFU, CR, Curcumin, Rapamycin, EGCG, NaCl, DTT and metal compounds along with control plates (without chemical agents). For, UV treatment, strains were spotted on to SC agar plates and exposed to 600µJ/cm^2^ UV radiation using UV crosslinker (CL-1000). For, 37°C treatment, strains were spotted on to SC agar plates and then the plate were incubated for 2-4 day at 37°C. All untreated and treated cells plates were incubated 30°C, the growth of the cells was recorded by scanning plates by HP scanner.

### Growth curve

The growth curve analysis was performed in liquid media. Overnight grown cells were inoculated and allowed to grow till the exponential phase. OD_600_ ∼ 0.2 of cells were seeded in duplicates in a 96-well cell culture plate (Tarsons-980040). Media without culture is taken as blank. The cells were allowed to grow continuously in the Eon™ Microplate spectrophotometer plate reader (Biotek) and OD_600_ values were measured at the regular interval of 30 minutes for 24 hours. Obtained readings from plate reader were processed in Microsoft excel 2013 and graph were plotted.

### Whole-cell protein extraction and western blotting

The whole-cell lysates were prepared by using the TCA precipitation method (51). For western blotting, whole cell extracts were resolved on 18% SDS-PAGE and proteins were transferred onto the 0.45µm nitrocellulose membrane (Bio-Rad, 1620115). The blots were blocked by using 2.5 % bovine serum albumin (Himedia, MB083) for 30 min, probed with the primary antibodies and secondary antibodies. The blots were visualized by IR-dye tagged secondary antibodies using the Odyssey infrared imaging system. Rad53 western blots were probed with HRP-conjugated anti-goat secondary antibody (RD systems) and developed using clarity western ECL substrate (Bio-Rad Laboratories, Inc, 170-5060). The anti-TBP western blotting was performed for the protein loading controls. List of antibodies is given in supplementary table 2. Blots were quantified using Image J software.

### Whole-cell protein extraction and western blotting for ubiquitination

Cells were grown till mid log phase and pelleted once with sterile MilliQ water. Lysate was prepared as previously described (26). Harvested treated and untreated cells were resuspended in Buffer B (50mM Tris-Cl 7.5 pH, 100 mM NaCl, 0.5% Triton-X-100, 0.1% deoxycholic acid and 1x complete mini protease inhibitor cocktail) and lysed using mechanical disruption. Cell lysates were collected and boiled for 5min in protein loading dye (0.1M Tris, 24% glycerol, 8% SDS, 0.2 mM dithiothreitol, 0.02% Coomassie blue G-250), proteins were separated using 4-20% gradient gel (Bio-Rad) and probed with anti-Ub antibody (Sigma-Aldrich). Ponceau-S staining of the blot was used as loading control. Details of antibodies are provided in the supplementary table 2.

### Total RNA extraction and gene expression analysis by real-time PCR

For gene expression studies, yeast cells were grown till the late log phase, harvested and total RNAs were prepared using rapid heat and freeze acidic phenol method. One microgram of RNA was used to synthesize cDNA using the iScript cDNA synthesis kit (Bio-Rad). Semi-quantitative PCR was performed using KAPA Taq PCR kit (KAPA biosystems) and amplified products were resolved on 1.2% Tris-EDTA-acetic acid buffer (TAE buffer). Standard protocols were used to perform Real-Time- quantitative PCR (Analytik jena qTOWER G) using TB Green Premix Ex Taq™ II (RR82LR, Takara). The *ACT1* gene was used as a control for the normalization of expression. Fold change relative to wild-type (untreated) cells is shown in the graph. The primers used in this study are listed in supplementary table 1.

### Chronological life span

Wild-type and H1246 strain were grown continuously for 10 days, aliquots of cells were taken every 2^nd^ day and serially diluted 10-fold. 3ul of serially diluted culture were spotted on SC agar plate. Plates were incubated at 30°C for 72 hours and the growth of cells were recorded by using HP scanner.

### Microscopy

To visualize lipid droplets inside yeast cells, BODIPY FL (493/503nm) 10mM stock was prepared in Dimethyl sulphoxide (DMSO). Overnight grown cells were inoculated at OD∼0.2 and allowed to grow at 30°C till OD_600_∼1. Untreated and MMS treated cell were further incubated for 2 hours in the shaker incubator at 30°C/200rpm. For staining, cells were harvested and washed twice with sterile 1X phosphate buffer saline (PBS) at 4°C/3000rpm/5min. cells were resuspended in 500µl of PBS and BODIPY was added to achieve concentration of 5µM. Cells were incubated for 20 min on rotamer and centrifuged at 3000rpm/5min. Cells were resuspended in PBS and visualized using Zeiss apotome DIC and e-GFP channels.

### Statistical Analysis

All the experiments were performed for at least three independent biological repeats or otherwise mentioned. The statistical analysis and significance were calculated using the student’s t-test relative to wild-type control. (\**p* < 0.05, \*\**p* < 0.001, \*\*\**p* < 0.0001; ns indicates non-significance). Obtained values were indicated as mean with standard error mean (SEM).

## Data availability

The authors declare that all data supporting the findings of the study is available within the manuscript and supplementary file.

## Supporting information

This article contains supporting information.

## Acknowledgement

We are grateful to Prof. Sten Stymne for gifting us WT (SCY62) and quadruple deletion mutant (H1246) and Prof. Susan Gasser for gifting us Mec1 and Mec1-S1991P antibodies. All the members of laboratory of chromatin biology are acknowledged for their suggestion.

## Author contribution

RS and RST designed the experiment. RS performed all the experiment. RS and RST analysed the results and wrote the manuscript.

## References

1. Jarc, E., and Petan, T. (2019) Lipid droplets and the management of cellular stress. The Yale journal of biology and medicine 92, 435

2. Müllner, H., and Daum, G. (2004) Dynamics of neutral lipid storage in yeast. Acta Biochimica Polonica 51, 323–347

3. Yang, H., Bard, M., Bruner, D. A., Gleeson, A., Deckelbaum, R. J., Aljinovic, G., Pohl, T. M., Rothstein, R., and Sturley, S. L. (1996) Sterol esterification in yeast: a two-gene process. Science 272, 1353–1356

4. Sandager, L., Gustavsson, M. H., Ståhl, U., Dahlqvist, A., Wiberg, E., Banas, A., Lenman, M., Ronne, H., and Stymne, S. (2002) Storage lipid synthesis is non-essential in yeast. Journal of Biological Chemistry 277, 6478–6482

5. Dubey, R., Stivala, C. E., Nguyen, H. Q., Goo, Y.-H., Paul, A., Carette, J. E., Trost, B. M., and Rohatgi, R. (2020) Lipid droplets can promote drug accumulation and activation. Nature chemical biology 16, 206–213

6. Barbosa, A. D., and Siniossoglou, S. (2020) New kid on the block: Lipid droplets in the nucleus. The FEBS journal 287, 4838–4843

7. Graef, M. (2018) Lipid dropletImediated lipid and protein homeostasis in budding yeast. FEBS letters 592, 1291–1303

8. Nguyen, T. B., Louie, S. M., Daniele, J. R., Tran, Q., Dillin, A., Zoncu, R., Nomura, D. K., and Olzmann, J. A. (2017) DGAT1-dependent lipid droplet biogenesis protects mitochondrial function during starvation-induced autophagy. Developmental cell 42, 9–21. e25

9. Yang, P.-L., Hsu, T.-H., Wang, C.-W., and Chen, R.-H. (2016) Lipid droplets maintain lipid homeostasis during anaphase for efficient cell separation in budding yeast. Molecular biology of the cell 27, 2368–2380

10. Stephenson, R. A., Thomalla, J. M., Chen, L., Kolkhof, P., White, R. P., Beller, M., and Welte, M. A. (2021) Sequestration to lipid droplets promotes histone availability by preventing turnover of excess histones. Development 148, dev199381

11. Ralhan, I., Chang, J., Moulton, M. J., Goodman, L. D., Lee, N. Y., Plummer, G., Pasolli, H. A., Matthies, D., Bellen, H. J., and Ioannou, M. S. (2023) Autolysosomal exocytosis of lipids protect neurons from ferroptosis. Journal of Cell Biology 222, e202207130

12. Shyu Jr, P., Wong, X. F. A., Crasta, K., and Thibault, G. (2018) Dropping in on lipid droplets: insights into cellular stress and cancer. Bioscience reports 38, BSR20180764

13. Hammerquist, A. M., Escorcia, W., and Curran, S. P. (2021) Maf1 regulates intracellular lipid homeostasis in response to DNA damage response activation. Molecular biology of the cell 32, 1086–1093

14. Ovejero, S., Kumanski, S., Soulet, C., Azarli, J., Pardo, B., Santt, O., Constantinou, A., Pasero, P., and MorielICarretero, M. (2023) A sterolIPI (4) P exchanger modulates the Tel1/ATM axis of the DNA damage response. The EMBO Journal 42, e112684

15. Zou, L., and Elledge, S. J. (2003) Sensing DNA damage through ATRIP recognition of RPA-ssDNA complexes. Science 300, 1542–1548

16. Cheng, Y.-H., Chuang, C.-N., Shen, H.-J., Lin, F.-M., and Wang, T.-F. (2013) Three distinct modes of Mec1/ATR and Tel1/ATM activation illustrate differential checkpoint targeting during budding yeast early meiosis. Molecular and cellular biology 33, 3365–3376

17. Wang, H., Liu, D., Wang, Y., Qin, J., and Elledge, S. J. (2001) Pds1 phosphorylation in response to DNA damage is essential for its DNA damage checkpoint function. Genes & development 15, 1361–1372

18. Mustafa, M., Habib, S., Imtiyaz, K., Tufail, N., Ahmad, R., Hamim, B., Abbas, K., Ahmad, W., Khan, S., and Rizvi, M. M. A. (2024) Characterization of structural, genotoxic, and immunological effects of methyl methanesulfonate (MMS) induced DNA modifications: Implications for inflammation-driven carcinogenesis. International Journal of Biological Macromolecules 268, 131743

19. Qiu, B., and Simon, M. C. (2016) BODIPY 493/503 staining of neutral lipid droplets for microscopy and quantification by flow cytometry. Bio-protocol 6, e1912–e1912

20. Hand, R. A., Jia, N., Bard, M., and Craven, R. J. (2003) Saccharomyces cerevisiae Dap1p, a novel DNA damage response protein related to the mammalian membrane-associated progesterone receptor. Eukaryotic cell 2, 306–317

21. Fu, Y. (2008) The regulatory network controlling DNA damage responses in Saccharomyces cerevisiae, University of Saskatchewan

22. Hustedt, N., Seeber, A., Sack, R., Tsai-Pflugfelder, M., Bhullar, B., Vlaming, H., van Leeuwen, F., Guénolé, A., van Attikum, H., and Srivas, R. (2015) Yeast PP4 interacts with ATR homolog Ddc2-Mec1 and regulates checkpoint signaling. Molecular cell 57, 273–289

23. Hurst, V., Challa, K., Jonas, F., Forey, R., Sack, R., Seebacher, J., Schmid, C. D., Barkai, N., Shimada, K., and Gasser, S. M. (2021) A regulatory phosphorylation site on Mec1 controls chromatin occupancy of RNA polymerases during replication stress. The EMBO Journal 40, e108439

24. Hanway, D., Chin, J. K., Xia, G., Oshiro, G., Winzeler, E. A., and Romesberg, F. E. (2002) Previously uncharacterized genes in the UV-and MMS-induced DNA damage response in yeast. Proceedings of the National Academy of Sciences 99, 10605–10610

25. Ghislain, M., Dohmen, R. J., Levy, F., and Varshavsky, A. (1996) Cdc48p interacts with Ufd3p, a WD repeat protein required for ubiquitinImediated proteolysis in Saccharomyces cerevisiae. The EMBO journal 15, 4884–4899

26. Lis, E. T., and Romesberg, F. E. (2006) Role of Doa1 in the Saccharomyces cerevisiae DNA damage response. Molecular and cellular biology 26, 4122–4133

27. Chung, N., Jenkins, G., Hannun, Y. A., Heitman, J., and Obeid, L. M. (2000) Sphingolipids signal heat stress-induced ubiquitin-dependent proteolysis. Journal of Biological Chemistry 275, 17229–17232

28. Zhang, J., Lei, Z., Huang, Z., Zhang, X., Zhou, Y., Luo, Z., Zeng, W., Su, J., Peng, C., and Chen, X. (2016) Epigallocatechin-3-gallate (EGCG) suppresses melanoma cell growth and metastasis by targeting TRAF6 activity. Oncotarget 7, 79557

29. Gault, C. R., Obeid, L. M., and Hannun, Y. A. (2010) An overview of sphingolipid metabolism: from synthesis to breakdown. Sphingolipids as signaling and regulatory molecules, 1-23

30. Rego, A., Trindade, D., Chaves, S. R., Manon, S., Costa, V., Sousa, M. J., and Côrte-Real, M. (2014) The yeast model system as a tool towards the understanding of apoptosis regulation by sphingolipids. FEMS yeast research 14, 160–178

31. Epstein, S., and Riezman, H. (2013) Sphingolipid signaling in yeast: potential implications for understanding disease. Front Biosci (Elite Ed) 5, 97–108

32. Dickson, R. C., and Lester, R. L. (2002) Sphingolipid functions in Saccharomyces cerevisiae. Biochimica et Biophysica Acta (BBA)-Molecular and Cell Biology of Lipids 1583, 13–25

33. Voynova, N. S., Vionnet, C., Ejsing, C. S., and Conzelmann, A. (2012) A novel pathway of ceramide metabolism in Saccharomyces cerevisiae. Biochemical Journal 447, 103–114

34. Al-razaq, A. (2024) Insights into the epigenetic role of H2A. J in radiation-induced premature senescence.

35. Isermann, A., Mann, C., and Rübe, C. E. (2020) Histone variant H2A. J marks persistent DNA damage and triggers the secretory phenotype in radiation-induced senescence. International journal of molecular sciences 21, 9130

36. Shokolenko, I. N., Wilson, G. L., and Alexeyev, M. F. (2013) Persistent damage induces mitochondrial DNA degradation. DNA repair 12, 488–499

37. Mirisola, M. G., and Longo, V. D. (2022) Yeast chronological lifespan: longevity regulatory genes and mechanisms. Cells 11, 1714

38. Duan, T., Taori, S., Bhargava, S., Lai, S., Zhong, C., Yomtoubian, S., Yuan, H., Wu, X., Zhang, P., and Huang, T. (2025) Nuclear cholesterol regulates nuclear size and DNA damage responses in cancer stem cells. *Neuro-Oncology*, noaf110

39. Cirigliano, A., Macone, A., Bianchi, M. M., Oliaro-Bosso, S., Balliano, G., Negri, R., and Rinaldi, T. (2019) Ergosterol reduction impairs mitochondrial DNA maintenance in S. cerevisiae. Biochimica et Biophysica Acta (BBA)-Molecular and Cell Biology of Lipids 1864, 290–303

40. Kovacs, M., Geltinger, F., Verwanger, T., Weiss, R., Richter, K., and Rinnerthaler, M. (2021) Lipid droplets protect aging mitochondria and thus promote lifespan in yeast cells. Frontiers in cell and developmental biology 9, 774985

41. Yu, L., Fan, J., Zhou, C., and Xu, C. (2021) Sterols are required for the coordinated assembly of lipid droplets in developing seeds. Nature Communications 12, 5598

42. Welte, M. A. (2015) Expanding roles for lipid droplets. Current biology 25, R470–R481

43. Chapman, K. D., Dyer, J. M., and Mullen, R. T. (2012) Biogenesis and functions of lipid droplets in plants: thematic review series: lipid droplet synthesis and metabolism: from yeast to man. Journal of lipid research 53, 215–226

44. Balakirev, M. Y., Mullally, J. E., Favier, A., Assard, N., Sulpice, E., Lindsey, D. F., Rulina, A. V., Gidrol, X., and Wilkinson, K. D. (2015) Wss1 metalloprotease partners with Cdc48/Doa1 in processing genotoxic SUMO conjugates. Elife 4, e06763

45. Geltinger, F., Tevini, J., Briza, P., Geiser, A., Bischof, J., Richter, K., Felder, T., and Rinnerthaler, M. (2020) The transfer of specific mitochondrial lipids and proteins to lipid droplets contributes to proteostasis upon stress and aging in the eukaryotic model system Saccharomyces cerevisiae. Geroscience 42, 19–38

46. Krick, R., Bremer, S., Welter, E., Schlotterhose, P., Muehe, Y., Eskelinen, E.-L., and Thumm, M. (2010) Cdc48/p97 and Shp1/p47 regulate autophagosome biogenesis in concert with ubiquitin-like Atg8. Journal of Cell Biology 190, 965–973

47. Bodnar, N. O., and Rapoport, T. A. (2017) Molecular mechanism of substrate processing by the Cdc48 ATPase complex. Cell 169, 722–735. e729

48. Lim, L., Jackson-Lewis, V., Wong, L., Shui, G., Goh, A., Kesavapany, S., Jenner, A., Fivaz, M., Przedborski, S., and Wenk, M. (2012) Lanosterol induces mitochondrial uncoupling and protects dopaminergic neurons from cell death in a model for Parkinson’s disease. Cell Death & Differentiation 19, 416–427

49. Dymond, J. S. (2013) Chapter twelveISaccharomyces cerevisiae growth media. Methods Enzymol 533, 191–204

50. Baker Brachmann, C., Davies, A., Cost, G. J., Caputo, E., Li, J., Hieter, P., and Boeke, J. D. (1998) Designer deletion strains derived from Saccharomyces cerevisiae S288C: a useful set of strains and plasmids for PCRImediated gene disruption and other applications. Yeast 14, 115–132

51. Singh, R., and Tomar, R. S. (2025) An Uncharacterized Domain Within the NITerminal Tail of Histone H3 Regulates the Transcription of FLO1 via Cyc8. Molecular Microbiology

