## Supporting file for "Lipid droplet-sphingolipid crosstalk regulates Doa1 activity and DNA damage response"

### Figure S1

## Ai

| | |
|---|---|
| WT (SCY62) | ***MATa*** *his3-11,15; leu2-3,112; ura3-1; trp1-1; can1-100 ADE2* |
| H1246 | ***MATα*** *are1Δ::HIS3 are2Δ::LEU2 dga1Δ::KanMX4 lro1Δ::TRP1 ADE2* |
| Q47 | ***Isogenic to*** **SCY62**<br>*are1Δ::HIS3;are2Δ::LEU2;lro1Δ::TRP1;dga1Δ::URA3* |

#### Aii

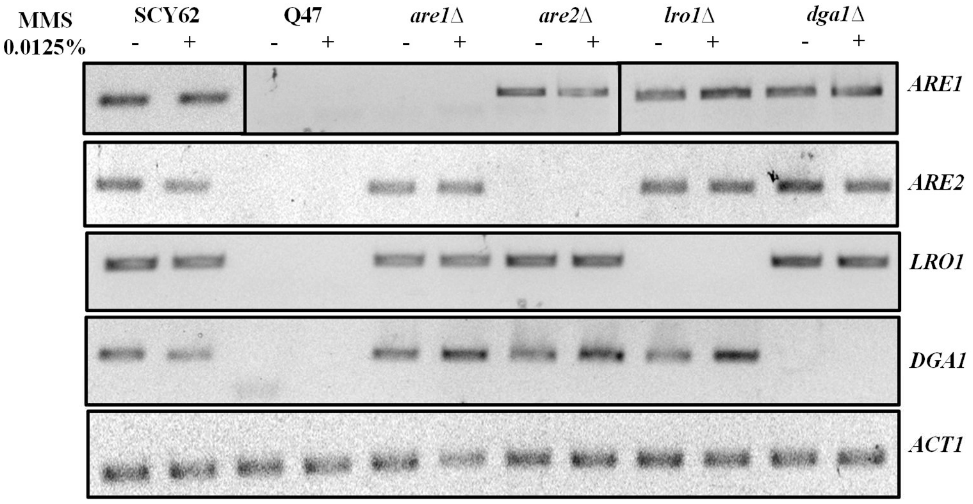


## B

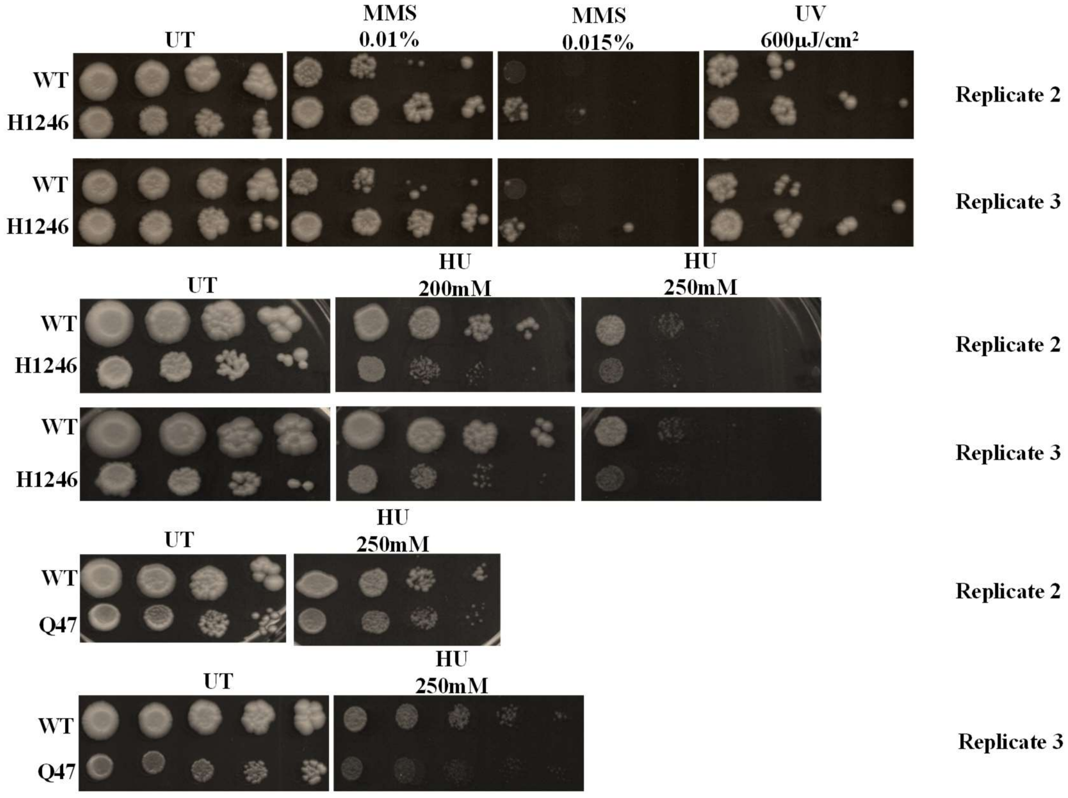

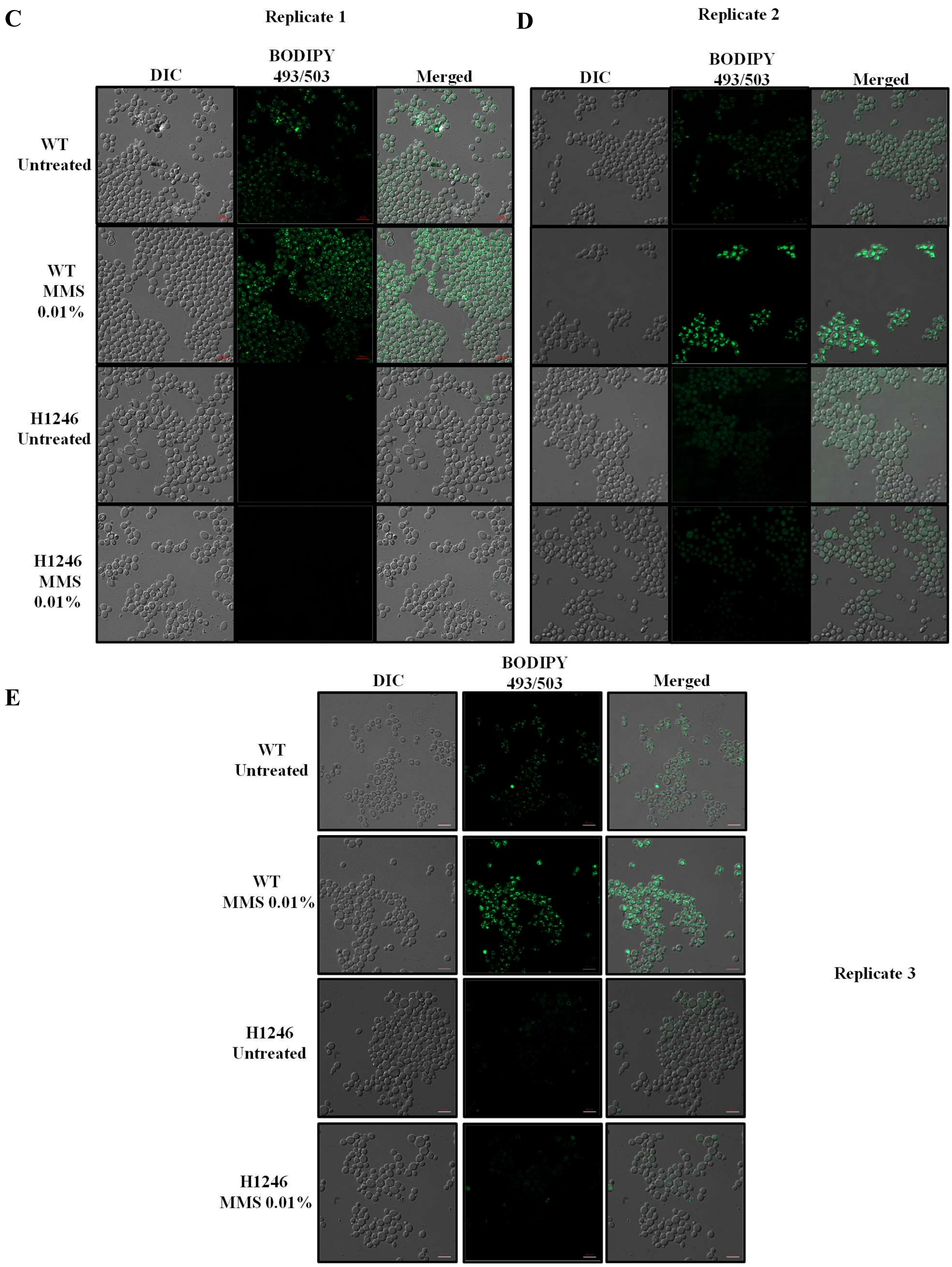
C
Replicate 1
DIC
BODIPY 493/503
Merged
WT Untreated
WT MMS 0.01%
H1246 Untreated
H1246 MMS 0.01%
D
Replicate 2
DIC
BODIPY 493/503
Merged
E
DIC
BODIPY 493/503
Merged
WT Untreated
WT MMS 0.01%
H1246 Untreated
H1246 MMS 0.01%
Replicate 3

**F**

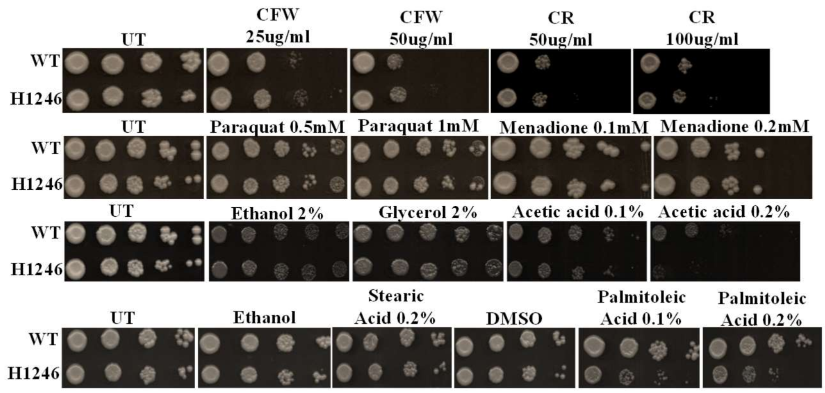


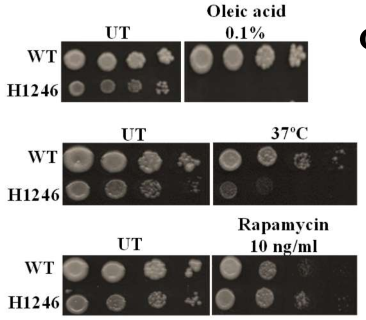


**G**

| S. No | Reagent | Sensitivity ration H1246/WT |
|---|---|---|
| 1 | **Acetic acid 0.2%** | 0.33 |
| 2 | **Palmitoleic acid 0.2%** | 0.5 |
| 3 | **Oleic acid 0.1%** | 0 |
| 4 | **Rapamycin** | 1.5 |
| 5 | **Curcumin** | 0.75 |
| 6 | **37°C** | 0.33 |
| 7 | **Tunicamycin 7.5nM** | 0.67 |

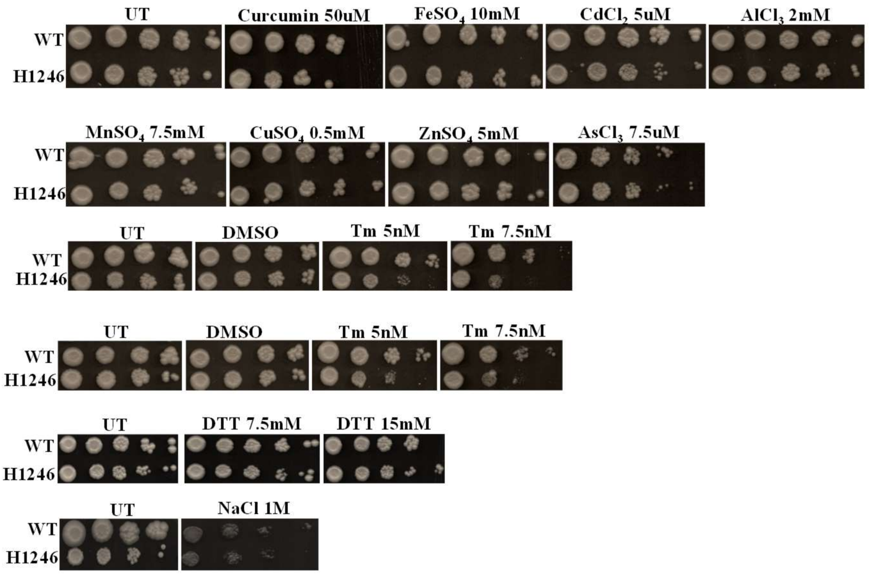


**Figure S1: (Ai)** Genotype of wild type (SCY62) and quadruple lipid droplets deficient mutant (H1246 and Q47). **(Aii)** Semi-quantitative PCR to confirm the gene deletions and testing expression of all the four genes of lipid biosynthesis pathway in untreated and MMS treated conditions. **(B)** Biological repeats of spot test assays of wild type along with H1246 and Q47 lipid droplets deficient mutants in absence and presence of different concentration of MMS (0.01% and 0.015%), hydroxy-urea (200 and 250mM) and UV exposure. **(C-E)** Fluorescence microscopic analysis of Bodipy (493/503nm) stained wild type and H1246 strains to monitor the cellular levels of lipid droplets upon MMS exposure. H1246, due to lack of lipid droplet biosynthesis genes, did not show any fluorescence upon staining with Bodipy, a dye specific to neutral lipid droplets. **(F)** Spot assay of wild type and H1246 lipid droplet deficient mutant cells upon treatment with different stress inducing agents. Cells were spotted in $10^4$ serial dilutions. **(G)** Growth sensitivity ration between the wild type and H1246 mutant under different stress inducing agents. Replicates depicts independent biological repeats of the respective experiments.

#### Figure S2

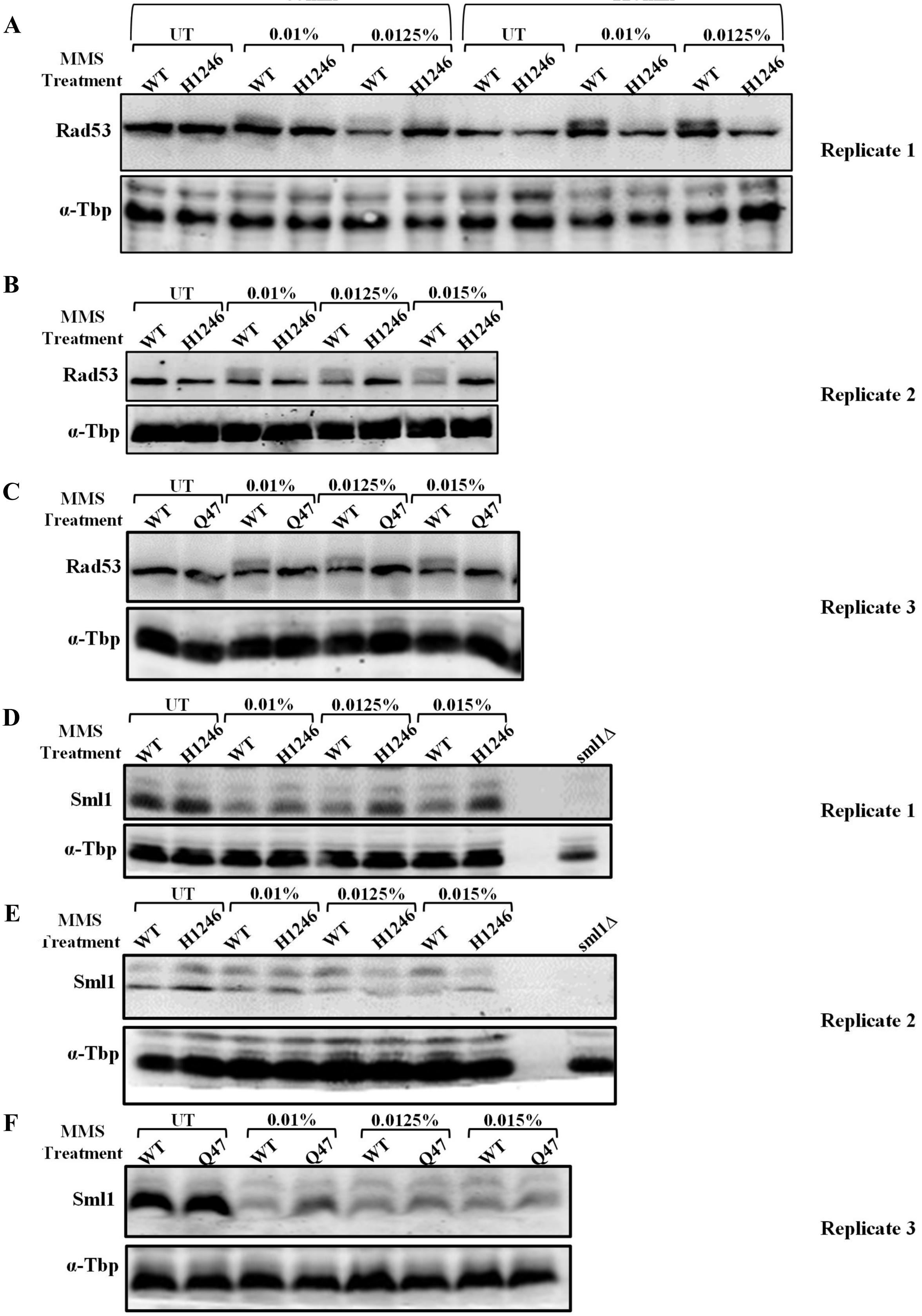

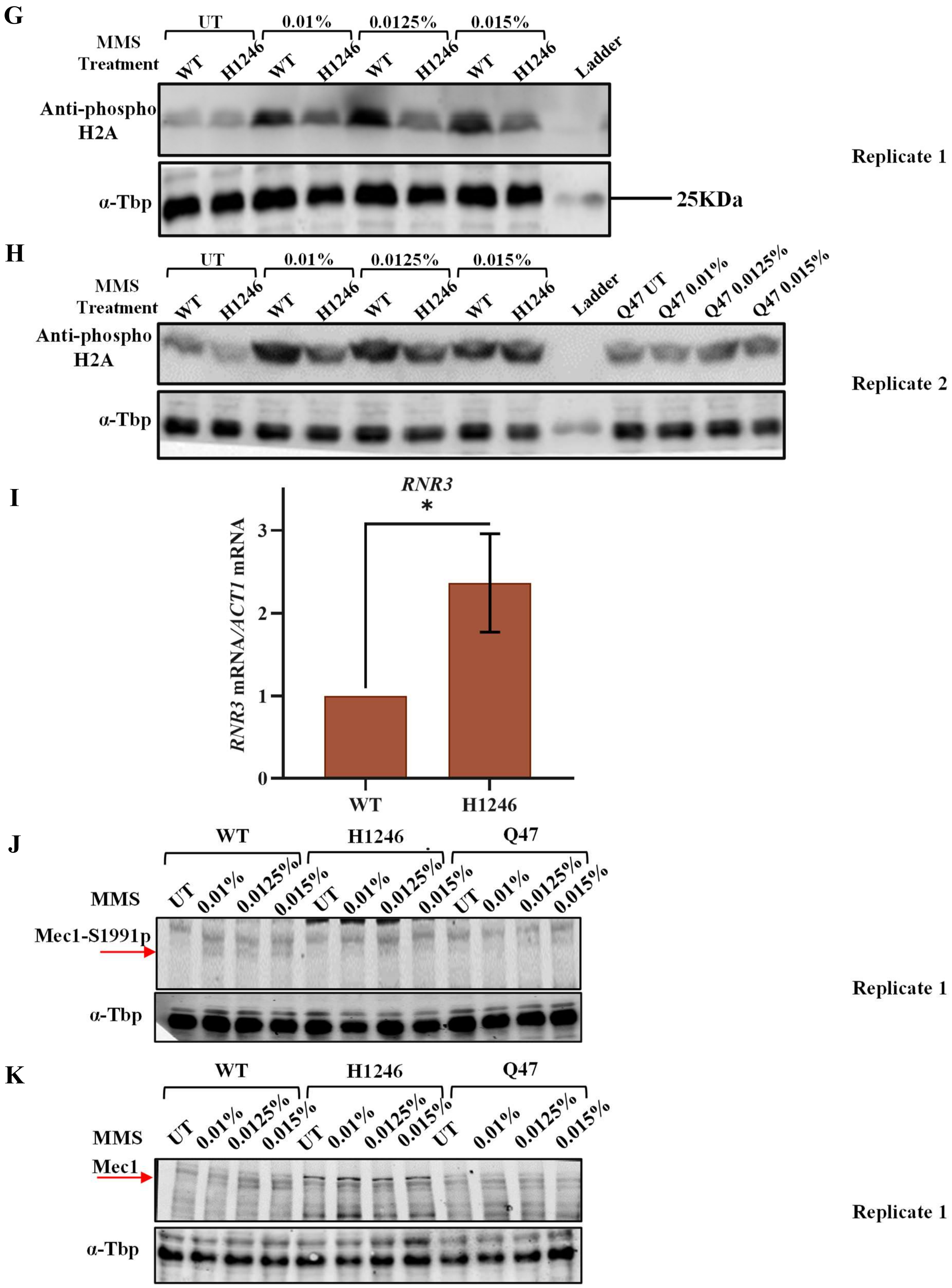
G
MMS Treatment
UT
0.01%
0.0125%
0.015%
WT
H1246
Ladder
Anti-phospho H2A
α-Tbp
25KDa
Replicate 1
H
MMS Treatment
UT
0.01%
0.0125%
0.015%
WT
H1246
Ladder
Q47 UT
Q47 0.01%
Q47 0.0125%
Q47 0.015%
Anti-phospho H2A
α-Tbp
Replicate 2
I
RNR3
*
RNR3 mRNA/ACT1 mRNA
0
1
2
3
WT
H1246
J
WT
H1246
Q47
MMS
UT
0.01%
0.0125%
0.015%
Mec1-S1991p
α-Tbp
Replicate 1
K
WT
H1246
Q47
MMS
UT
0.01%
0.0125%
0.015%
Mec1
α-Tbp
Replicate 1

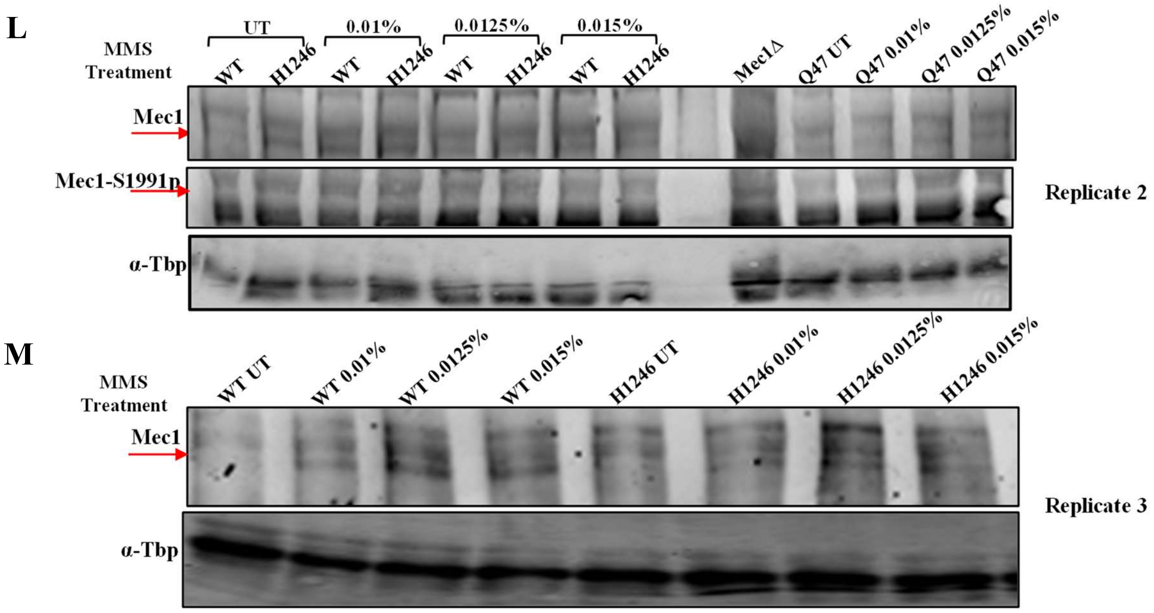


**Figure S2: (A)** Immunoblot to measure Rad53 phosphorylation levels in wild type and H1246 lipid droplet deficient mutant cells upon treatment with different concentrations of MMS ranging from 0.01% - 0.0125% for 60 and 120 minutes. Whole cell extracts were resolved on 12% SDS-PAGE. The western blot with anti-Tbp is used for loading control. **(B)** Immunoblotting of Rad53 to measure Rad53 phosphorylation levels in wild type and H1246 mutant strain upon 0.01%, 0.0125% and 0.015% MMS treatments for 120 minutes. **(C)** Immunoblotting to measure Rad53 phosphorylation levels in wild type and Q47 lipid droplets deficient mutant strain upon MMS treatments (0.01%, 0.0125% and 0.015%) for 120 minutes. **(D-E)** Biological repeats of immunoblotting of Sml1 to examine degradation of Sml1 protein in response to MMS treatment in wild type and H1246 mutant. **(F)** Immunoblotting of Sml1 to measure Sml1 degradation in response to MMS treatment in wild type and Q47 lipid droplets deficient mutant. sml1Δ mutant strain was taken as a negative control for Sml1 immunoblotting. Whole cell extracts were resolved through 18% SDS-PAGE. The α-Tbp western blotting was used for the protein loading control. **(G-H)** Biological repeats of western blots to measure histone H2A phosphorylation in wild type and lipid droplets deficient mutants (H1246 and Q47) upon MMS treatment. Whole cell protein extracts were resolved on 18% SDS-polyacrylamide gel. α-Tbp western blotting was performed for protein loading control. **(I)** RT-PCR analysis to measure basal expression level of *RNR3* in wild type and H1246 mutant strain. PCR data were processed in Microsoft-excel and graph was plotted. **(J-M)** Western blotting to measure levels of Mec1 and phosphorylation at Mec1-S1991 site. Whole cell extracts were resolved on 4-20% gradient gel. Western using anti-Tbp antibody was conducted for the protein loading control.

### Figure S3

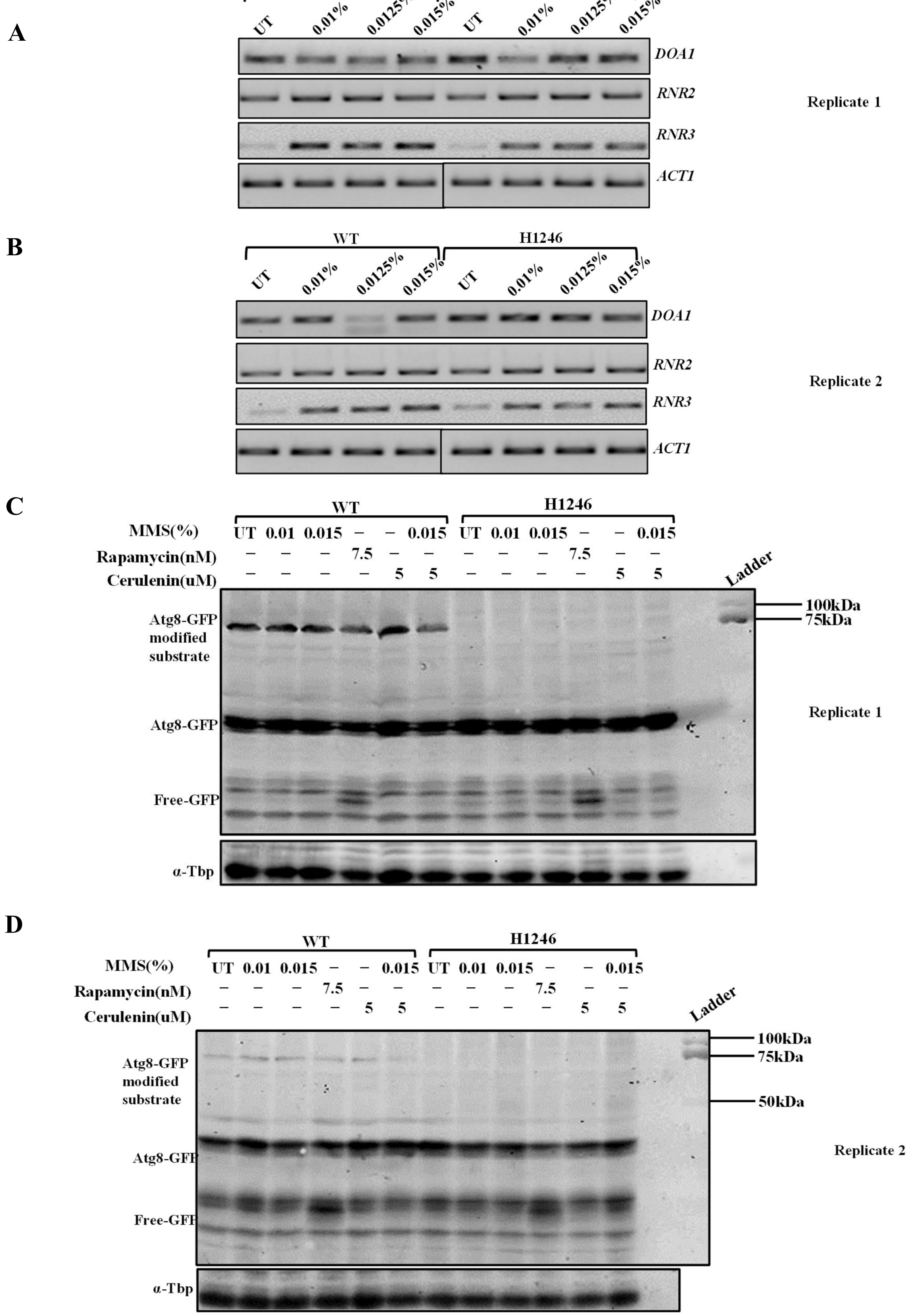
A
WT
H1246
UT
0.01%
0.0125%
0.015%
DOA1
RNR2
RNR3
ACT1
Replicate 1
B
WT
H1246
UT
0.01%
0.0125%
0.015%
DOA1
RNR2
RNR3
ACT1
Replicate 2
C
WT
H1246
MMS(%) UT 0.01 0.015 – – 0.015 UT 0.01 0.015 – – 0.015
Rapamycin(nM) – – – 7.5 – – – – – 7.5 – –
Cerulenin(uM) – – – – 5 5 – – – – 5 5
Ladder
Atg8-GFP modified substrate
Atg8-GFP
Free-GFP
α-Tbp
100kDa
75kDa
Replicate 1
D
WT
H1246
MMS(%) UT 0.01 0.015 – – 0.015 UT 0.01 0.015 – – 0.015
Rapamycin(nM) – – – 7.5 – – – – – 7.5 – –
Cerulenin(uM) – – – – 5 5 – – – – 5 5
Ladder
Atg8-GFP modified substrate
Atg8-GFP
Free-GFP
α-Tbp
100kDa
75kDa
50kDa
Replicate 2

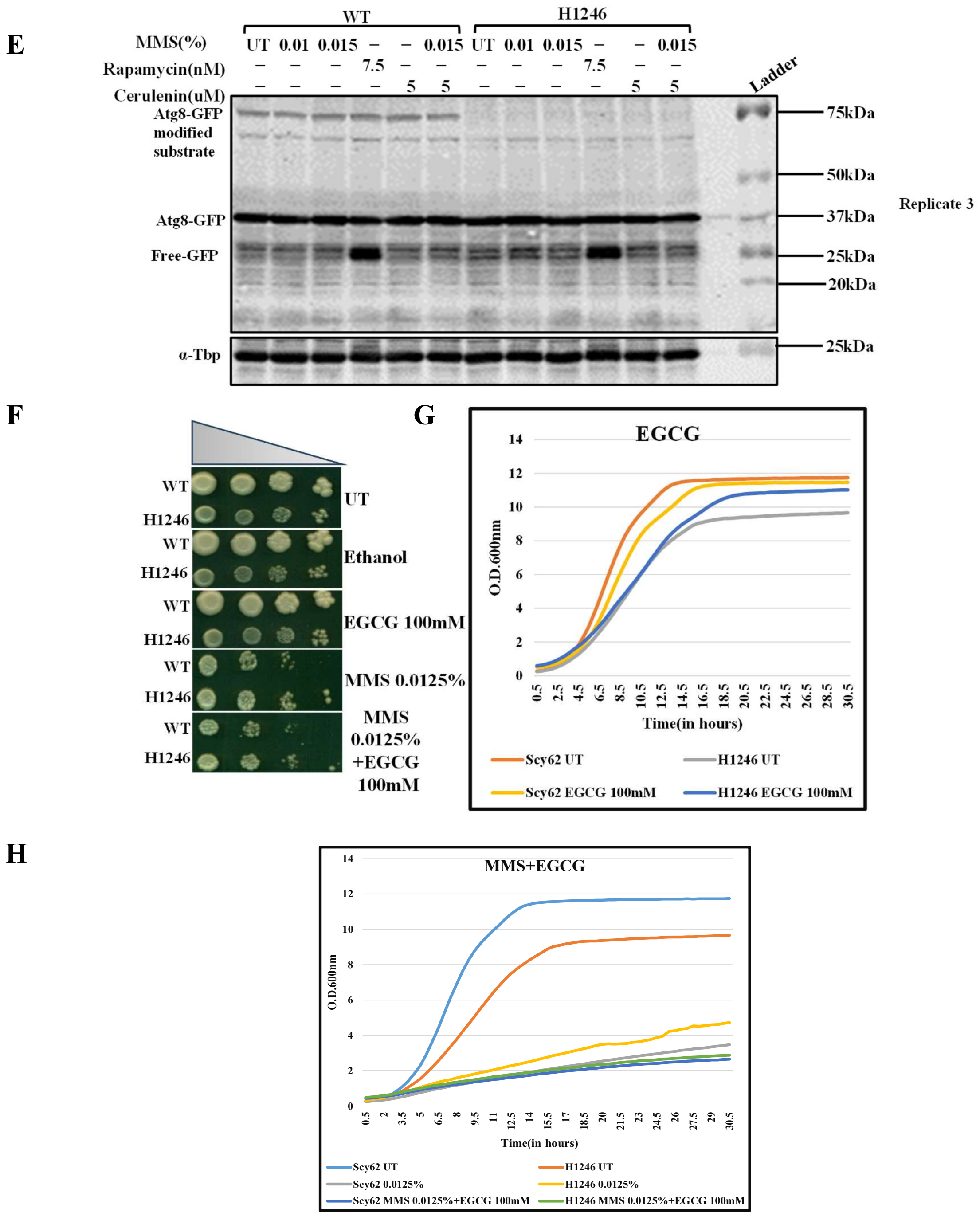


**Figure 3 (A, B)** Biological repeats of semi-quantitative-PCR to measure expression of *DOA1, RNR2* and *RNR3* genes in wild type and H1246 lipid droplet deficient mutant in untreated and upon MMS treatments for 2 hours. **(C)** GFP immunoblotting by using anti-GFP antibody to measure GFP-Atg8 and free GFP levels in wild type and H1246 mutant. Band of ~75KDa is most likely a Atg8 modified substrate which is absent in H1246 mutant probably got degraded due to hyperactivity of Doa1 and Ccd48 proteasome. Rapamycin treatment to induce autophagy, ATG8 substrate does not get affected upon rapamycin treatment in wild type cells. **(D, E)** Biological repeats of GFP-Atg8 western blots. **(F)** Spot assay to examine the effect of 100mM EGCG (E3 ligase inhibitor) on the growth of wild type and H1246 mutant strain in untreated and MMS treatment condition. **(G, H)** Growth curve analysis to examine the effect of EGCG on wild type and H1246 mutant in untreated and under MMS treatment.

#### Figure S4

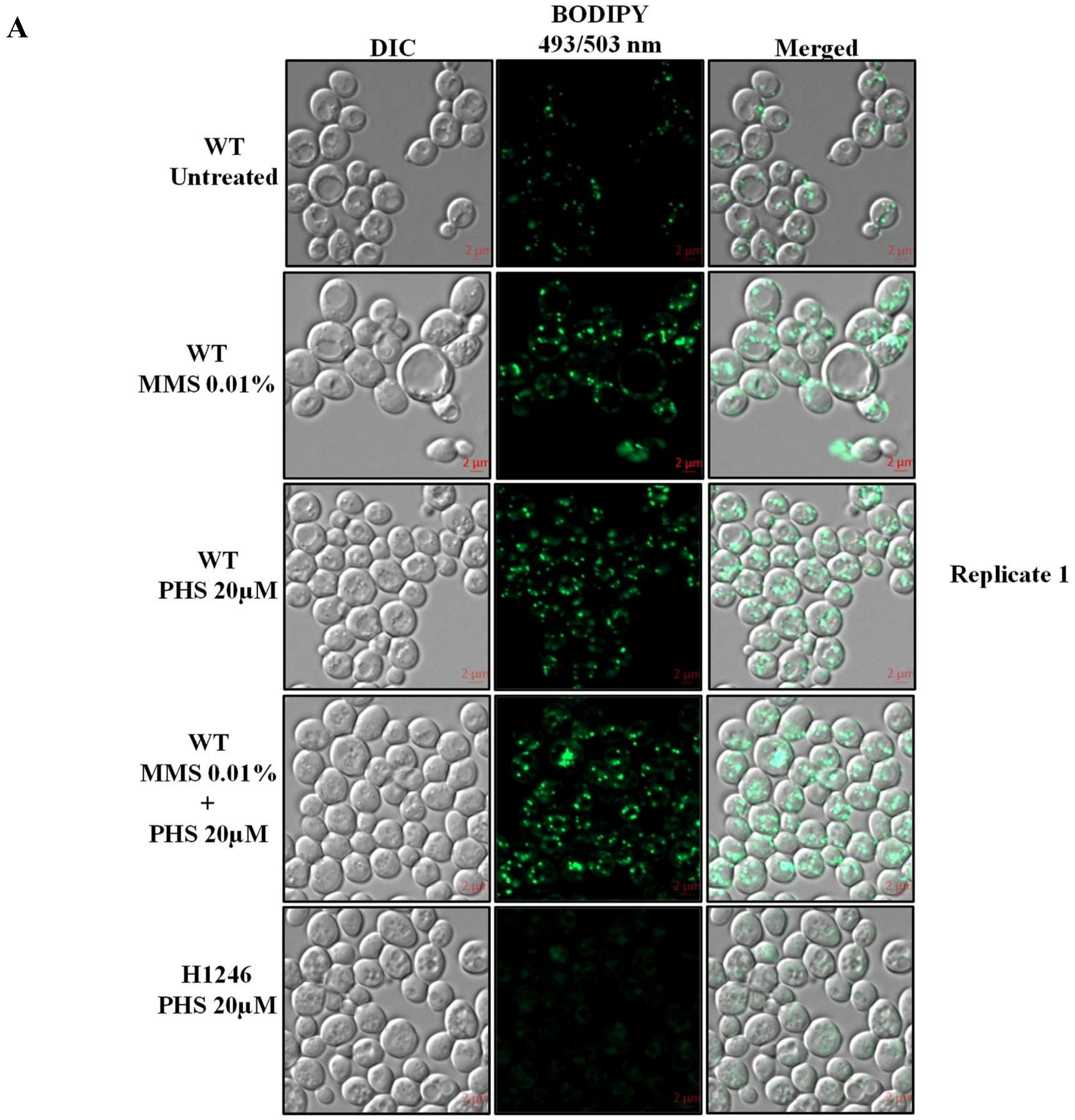
A
DIC
BODIPY 493/503 nm
Merged
WT Untreated
WT MMS 0.01%
WT PHS 20μM
WT MMS 0.01% + PHS 20μM
H1246 PHS 20μM
Replicate 1

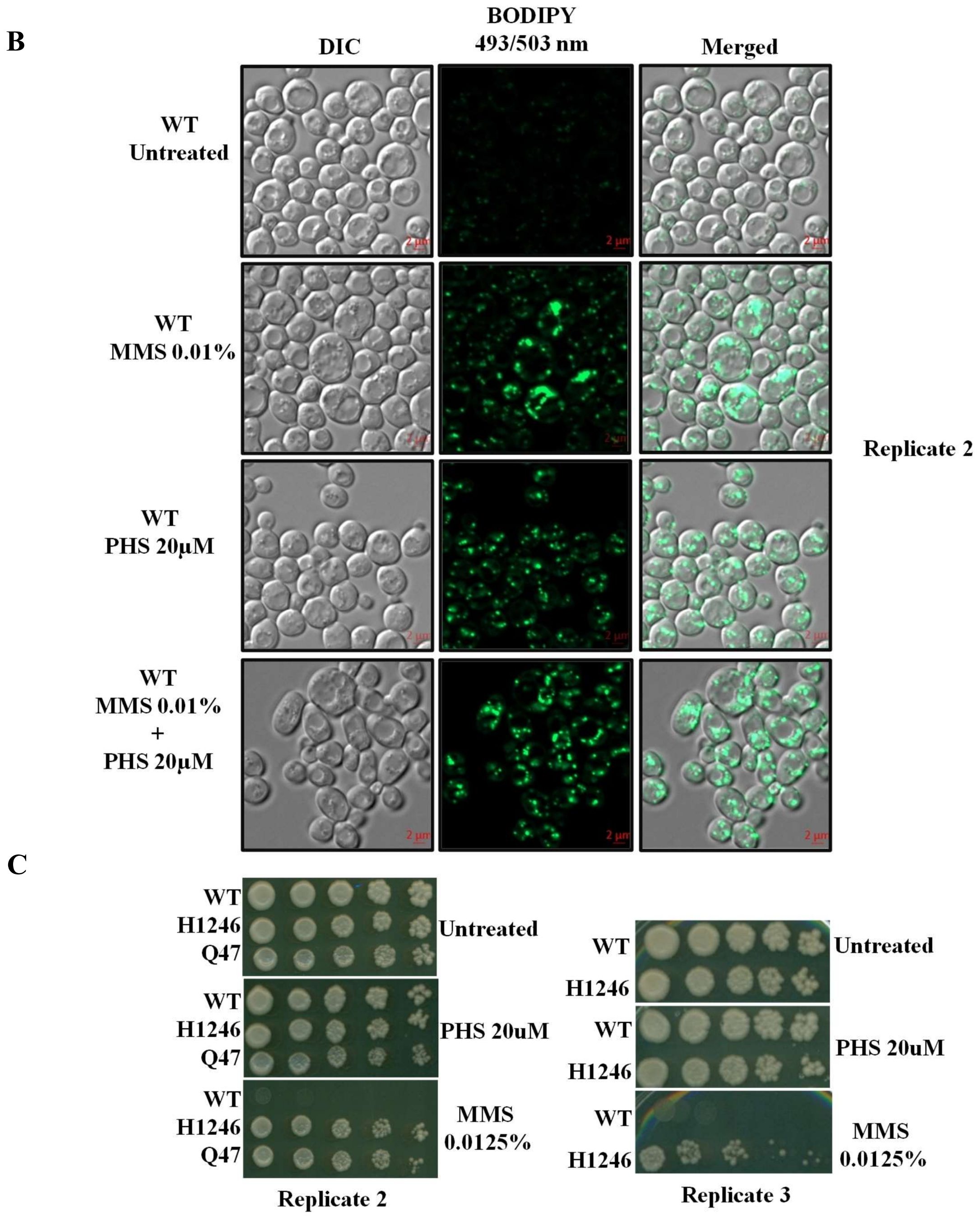


**Figure 4: (A-B)** Fluorescence microscopic analysis of Bodipy stained cells to examine the effect of PHS treatment on lipid droplet formation. Cells were stained with Bodipy post 20uM PHS treatment for 2hrs. **(C)** Repeats of spot assay to test the PHS effect on growth of wild type and H1246 mutant. PHS does not induce growth defect but increases lipid droplets count which may be due to neutralization of excess cytosolic PHS and deposition in the lipid droplets. Droplets count increases in wild type to store excess PHS but no change in the mutant.

#### Figure S5

A

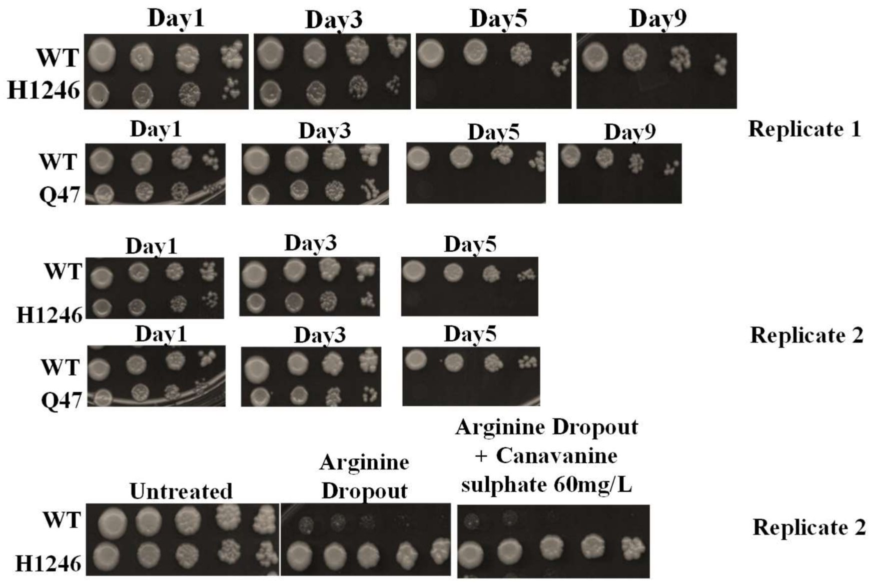


**Figure 5: (A-B)** Biological replicates of spot assay to determine chronological life span (CLS) of wild type and lipid droplets deficient mutants; H1246 and Q47. Biological replicate-3 was performed to measure CLS only till day 5 because H1246 grows only till day 3. The growth of wild type cells was not affected till day 9. **(C)** Biological replicate of spot assays to test the growth of wild type and H1246 mutant cells on canavanine sulphate (60mg/L, a toxic arginine analogue). H1246 mutant cells do not respond to DDR (double strand break), mutation in arginine permease could grow on canavanine containing media but wild type cells show hypersensitivity to canavanine sulphate.

#### Figure S6

A B C

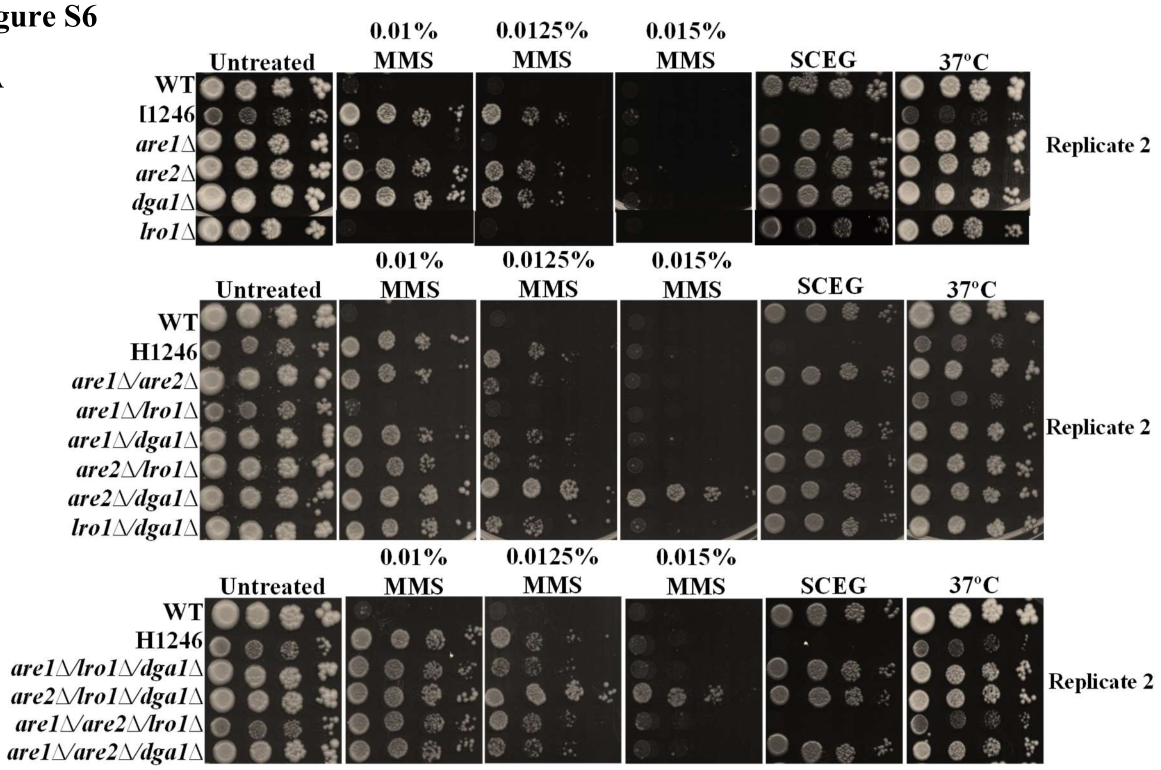

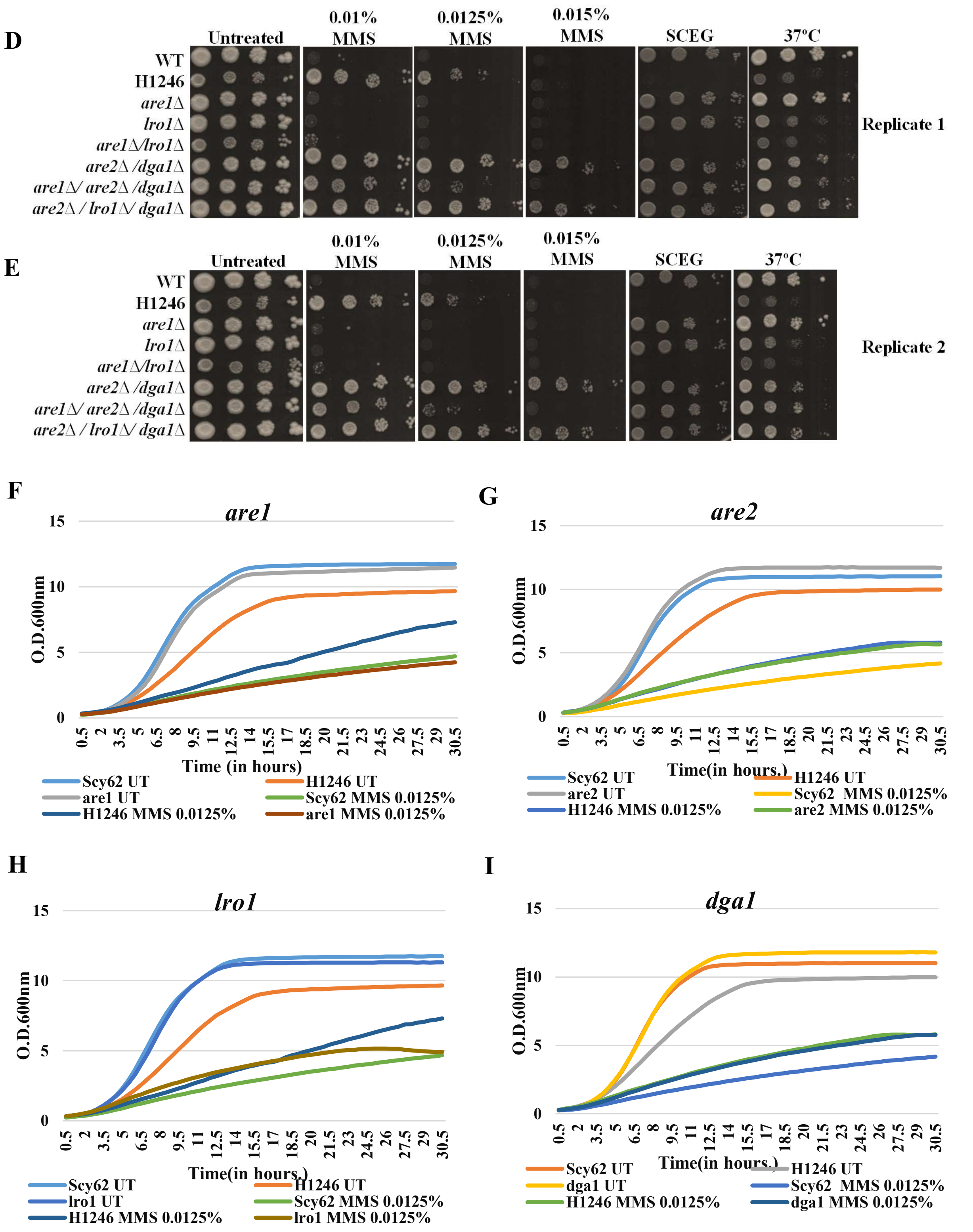


**Figure 6: (A-C)** Spot assay to examine the growth phenotype of single, double and triple deletion mutants of lipid biosynthesis gene in untreated and in MMS containing SC media, SCEG media and at 37ºC. Red text depicts MMS resistant mutant and yellow text depicts sensitive mutants. **(D-E)** Biological replicates of spot assays to assess the growth of selected mutant strains which are more sensitivity or resistance. **(F-I)** Growth kinetics of single deletion mutants are1Δ, are2Δ, lro1Δ and dga1Δ, respectively. Kinetics show that are1Δ and lro1Δ show

similar growth phenotype as wild type cells whereas are2Δ and dga1Δ show growth phenotype same as quadruple mutant, H1246 upon MMS treatment.

#### Figure S7

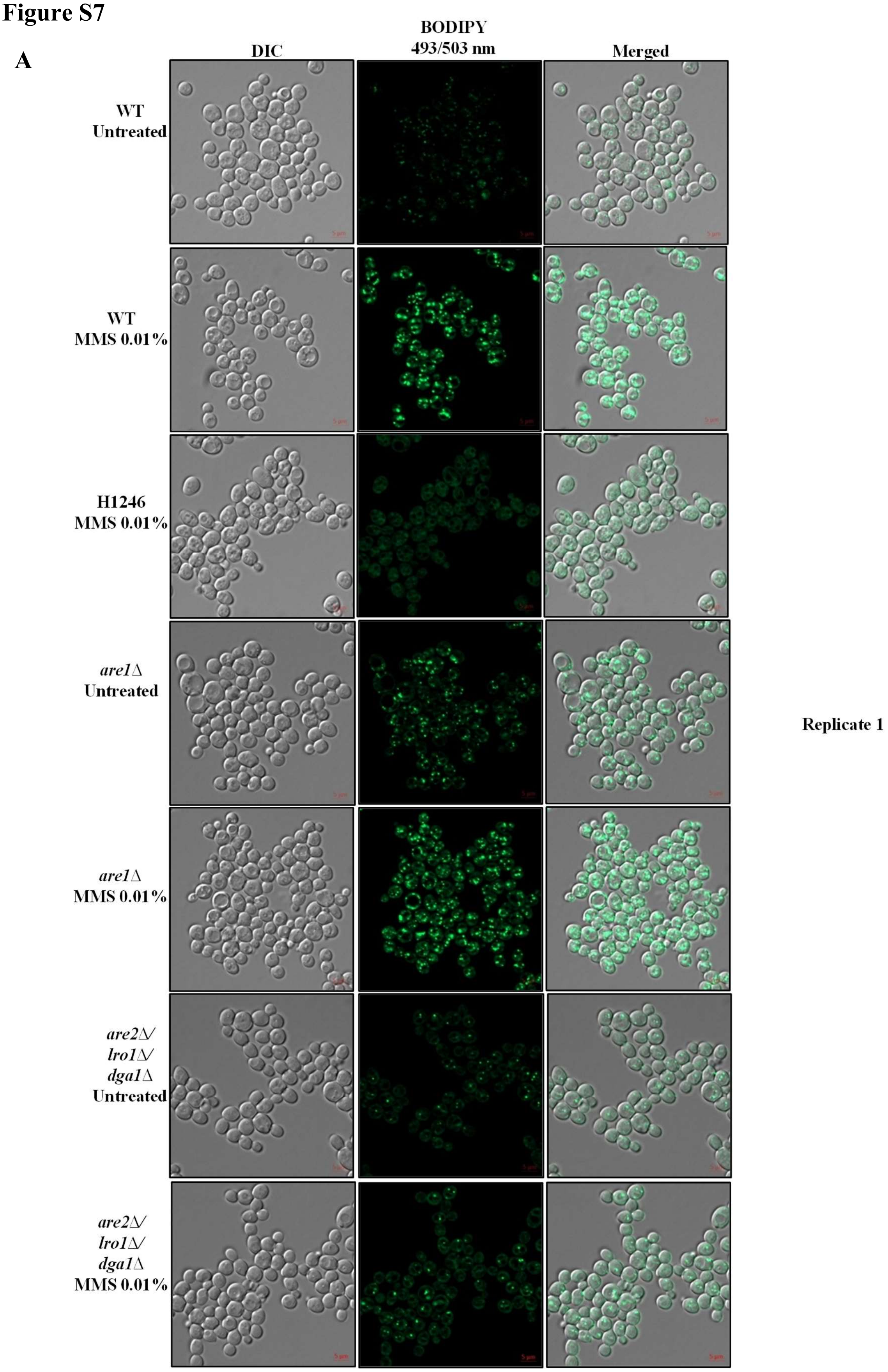

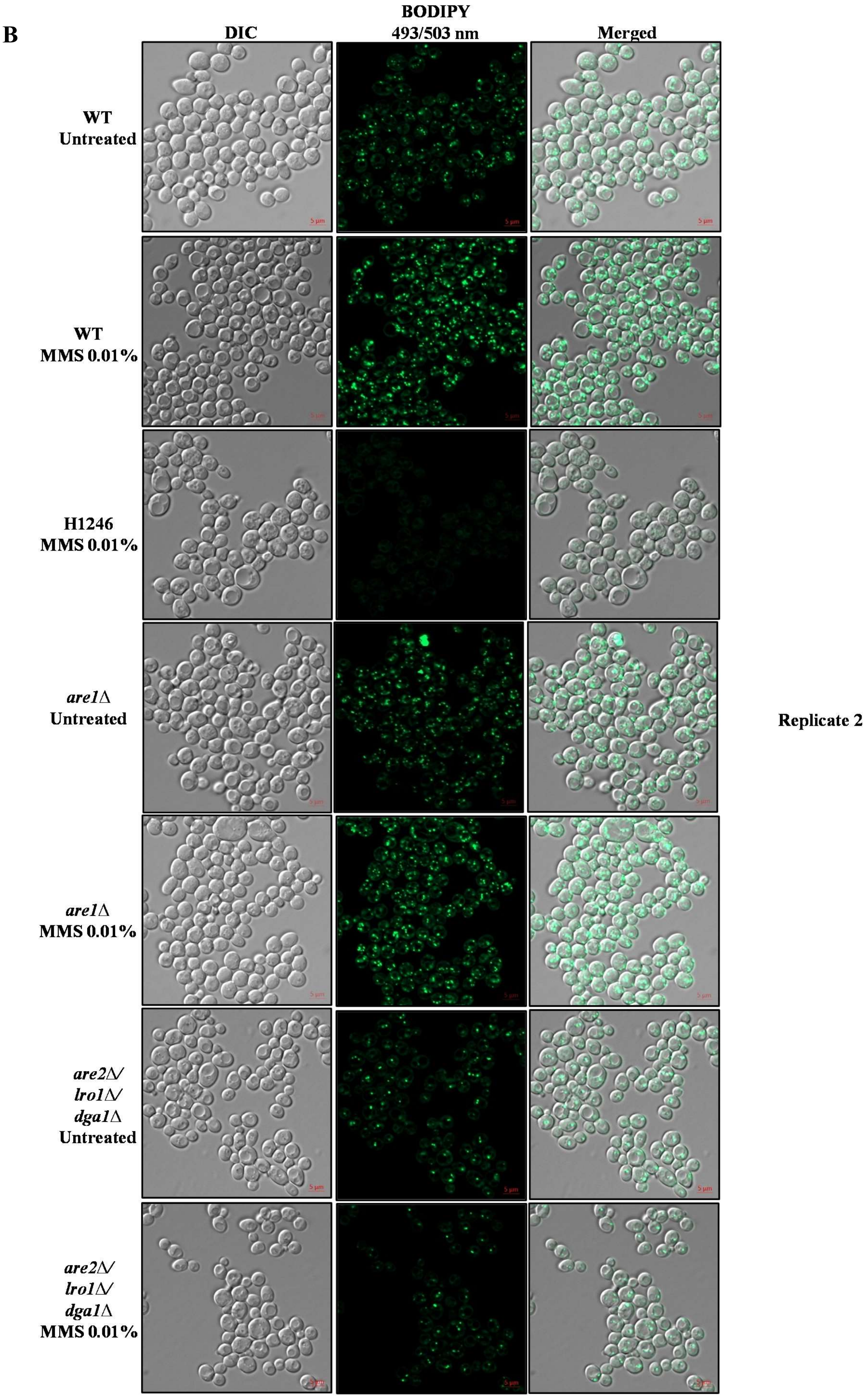
B
DIC
BODIPY
493/503 nm
Merged
WT
Untreated
WT
MMS 0.01%
H1246
MMS 0.01%
*are1*Δ
Untreated
*are1*Δ
MMS 0.01%
*are2*Δ/
*lro1*Δ/
*dga1*Δ
Untreated
*are2*Δ/
*lro1*Δ/
*dga1*Δ
MMS 0.01%
Replicate 2

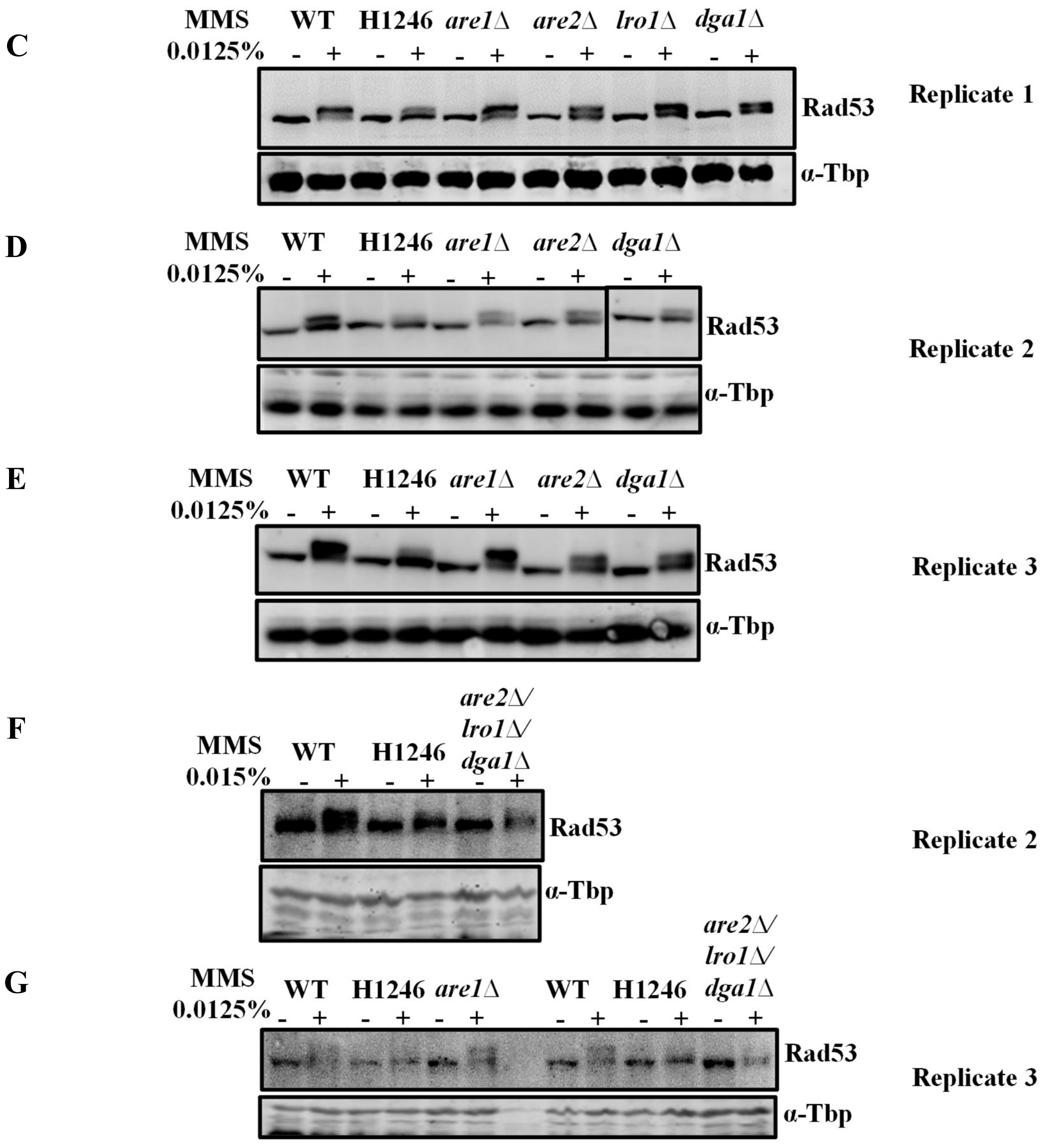


**Figure S7: (A-B)** Fluorescence microscopic analysis by using a lipid droplets specific dye, Bodipy to compare the lipid droplets formation in wild type, are1Δ and triple deletion mutant, are2Δ/lro1Δ/dga1Δ in untreated and upon MMS treatment condition. Triple deletion mutant show very low level of LD count even after MMS treatment. **(C-E)** Biological replicates of immunoblotting to measure Rad53 phosphorylation in are1Δ mutant and compared with the wild type cells in untreated and upon MMS treatment. We observed much higher density of lipid droplets and more Rad53 phosphorylation in are1Δ strain in comparison to wild type cells. **(F, H)** Rad53 immunoblotting to monitor phosphorylation status in are2Δ/lro1Δ/dga1Δ in untreated and MMS treatment. In biological replicate 3, are1Δ deletion was taken to compare the Rad53 phosphorylation between are1Δ and are2Δ/lro1Δ/dga1Δ mutants along with wild type cells in untreated and upon 0.0125% MMS treatment for 2 hrs.

#### Supplementary table 1: List of primers

| S. No. | Name | Primer sequence |
|---|---|---|
| 1 | *ARE1 (F)* | AACAAACGGCATTCGGTCAC |

| | | |
|---|---|---|
| 2 | *ARE1 (R)* | AGGTACTTTTGCATGTGGGC |
| 3 | *ARE2 (F)* | CACCTCAGAAACGGTGGTCA |
| 4 | *ARE2 (R)* | CTACGGATGCCACAGTGAAC |
| 5 | *LRO1 (F)* | CCTGGTGTCATTTCTACGGGA |
| 6 | *LRO1 (R)* | GCACGTAGCGTAAAGTTCGG |
| 7 | *DGA1 (F)* | GCGTTTGCAACAGAAGGTTG |
| 8 | *DGA1 (R)* | CTAGCGCCACCAACAACAAT |
| 9 | *DOA1 (F)* | GGATGTGGTAGCTGTGGATGA |
| 10 | *DOA1 (R)* | ACAAGGGCACACCGTTGAT |
| 11 | *SUR2 (F)* | TATGCACTATGCCAAGGCCC |
| 12 | *SUR2 (R)* | TGTTCCTCTTGGCAACCTCTTC |
| 13 | *YPC1 (F)* | GGCGTTTGGGGAGAAACAACT |
| 14 | *YPC1 (R)* | ACTCCGACCAAACCGTACCC |
| 15 | *YDC1 (F)* | AGGTTATTGGGGCAAGCCAAC |
| 16 | *YDC1 (R)* | TACCAACCAGCGAGAACCCC |
| 17 | *YSR3 (F)* | CCTGTGTGGCTTGGATACCG |
| 18 | *YSR3 (R)* | ACCGCAGTAGCGTTAGCAG |
| 19 | *LCB3 (F)* | GCTCCAAGCTCCCATACAGC |
| 20 | *LCB3 (R)* | CCAATGAGCCCACCGCTTA |
| 21 | *LCB4 (F)* | TCCTGACAAAAGCAAGGCCA |
| 22 | *LCB4 (R)* | CGCATCCACTCTGTCGGG |
| 23 | *DPL1 (F)* | GTATGCCCTTCATCATCGTGGA |
| 24 | *DPL1 (R)* | GAGCGGAACCGACCAGTAAA |
| 25 | *AUR1 (F)* | GCCCCATTTGTCGTTGCTG |

| | | |
|---|---|---|
| 26 | *AUR1 (R)* | GCTAATCCACCAGGCGAGC |
| 27 | *IPT1 (F)* | CCATCTGCTTGTTCCAATGGC |
| 28 | *IPT1 (R)* | TGGCGGAGTGTAGTGAAGGA |
| 29 | *CHS1 (F)* | GAAGCTGTGAGAGGCCAACC |
| 30 | *CHS1 (R)* | ACCCTCTCCGGTACATCTGTT |
| 31 | *RNR2 (F)* | CCGCTTCTGACGGTATTGTT |
| 32 | *RNR2 (R)* | TCTGGGATGGTGTGAATGG |
| 33 | *RNR3 (F)* | TCGGATTCAAGACACTGGAC |
| 34 | *RNR3 (R)* | TAAAGTTGGGGAAGCGTGAG |
| 35 | *COX1 (F)* | CACCACTAATTGAAAACCTGTCTG |
| 36 | *COX1 (R)* | GATTTATCGTATGCTCATTTCCAA |
| 37 | *COX2 (F)* | GCTGATGTTATTCATGATTTTGCT |
| 38 | *COX2 (R)* | TGGCATATTTGCATGACCTG |
| 39 | *ACT1 (F)* | TCGTCGGTAGACCAAGACAC |
| 40 | *ACT1 (R)* | TTCTTCTGGGGCAACTCTCA |
| 39 | *ARE1 K.O. (F)* | AGCACGGCTTGCAGCAAGAGCGCCAAAACA GATTGCAAGACTGTGCGGTATTTCACACCG |
| 40 | *ARE1 K.O. (R)* | ATCAAGGGCTTGCGAGGGACACACGTGGTATGGTGGCAGT AGATTGTACTGAGAGTGCAC |
| 41 | *ARE2 K.O. (F)* | ACACATTACGTTAGCAAAAGCAACAATAACAAACACAACC CTGTGCGGTATTTCACACCG |
| 42 | *ARE2 K.O. (R)* | ATAAAGATTTAATAGCTCCACAGAACAGTTGCAGGATGCC AGATTGTACTGAGAGTGCAC |
| 43 | *LRO1 K.O. (F)* | AGGTTCTCTACCAACGAATTCGGCGACAATCGAGTAAAAA CTGTGCGGTATTTCACACCG |
| 44 | *LRO1 K.O. (R)* | GAAATAATACACGGATGGATAGTGAGTCAATGTCGGTCAT AGATTGTACTGAGAGTGCAC |

| | | |
|---|---|---|
| 45 | *DGA1 K.O. (F)* | TAAGGAAACGCAGAGGCATACAGTTTGAACAGTCACATAA CTGTGCGGTATTTCACACCG |
| 46 | *DGA1 K.O. (R)* | TTTATTCTAACATATTTTGTGTTTTCCAATGAATTCATTA AGATTGTACTGAGAGTGCAC |
| 47 | *DOA1 K.O. (F)* | TCATGTGTGATAGTAAGGTGTAGAGCAGCAGATTTGGAGT CTGTGCGGTATTTCACACCG |
| 48 | *DOA1 K.O. (R)* | ATCTAGACATTATGTGTTTTATATGATTGCTGTAAAAGTAA GATTGTACTGAGAGTGCAC |

**Supplementary table 2: List of antibodies**

| S. No. | Antibody | Catalogue No. | Manufacturer |
|---|---|---|---|
| 1 | α-Tbp | - | In house |
| 2 | Rad53 | SC-6749 | Santa Cruz Biotechnology |
| 3 | Sml1 | AS10 847 | Agri sera |
| 4 | Anti-phospho Histone H2A(Serine 129) | 07-745 | Merck |
| 5 | Anti-Ubiquitin | SAB4503053 | Merck |
| 6 | Anti-GFP | G1544 | Sigma-Aldrich |
| 7 | Mec1 | - | Gifted By Prof. S.M. Gasser |
| 8 | Mec1-S1991P | - | Gifted By Prof. S.M. Gasser |
| 9 | Goat anti-Rabbit IgG (H+L) Secondary Antibody, Alexa Fluor™ Plus 680 | A32734 | Thermo Fisher scientific |
| 10 | Goat anti-Rabbit IgG (H+L) Secondary Antibody, Alexa Fluor™ Plus 800 | A32735 | Thermo Fisher scientific |

**Supplementary table 3: List of strains**

| **Strain** | **Genotype** | **Source** |
|---|---|---|
| WT (SCY62) | *MATa his3-11,15; leu2-3,112; ura3-1; trp1-1; can1-100 ADE2* | Gifted by Prof. Stymne |
| H1246 | *MATα are1Δ::HIS3 are2Δ::LEU2 dga1Δ::KanMX4 lro1Δ::TRP1 ADE2* | Gifted by Prof. Stymne |
| Q47 | *Isogenic to* Scy62 *are1Δ::HIS3;are2Δ::LEU2;lro1Δ::TRP1;dga1Δ::URA3* | This study |
| *are1Δ* | *Isogenic to* Scy62 *are1Δ::HIS3* | This study |
| *are2Δ* | *Isogenic to* Scy62 *are2Δ::HIS3* | This study |

| | | |
|---|---|---|
| *lro1Δ* | *Isogenic to* Scy62 *lro1Δ::LEU2* | This study |
| *dga1Δ* | *Isogenic to* Scy62 *dga1Δ::TRP1* | This study |
| *are1Δ/are2Δ* | *Isogenic to* Scy62 *are1Δ::HIS3;are2Δ::TRP1* | This study |
| *are1Δ/lro1Δ* | *Isogenic to* Scy62 *are1Δ::HIS3;lro1Δ::TRP1* | This study |
| *are1Δ/dga1Δ* | *Isogenic to* Scy62 *are1Δ::HIS3;dga1Δ::URA3* | This study |
| *are2Δ/lro1Δ* | *Isogenic to* Scy62 *are2Δ::HIS3;lro1Δ::TRP1* | This study |
| *are2Δ/dga1Δ* | *Isogenic to* Scy62 *are2Δ::HIS3;dga1Δ::URA3* | This study |
| *lro1Δ/dga1Δ* | *Isogenic to* Scy62 *lro1Δ::TRP1;dga1Δ::URA3* | This study |
| *are1Δ/are2Δ/lro1Δ* | *Isogenic to* Scy62 *are1Δ::HIS3;are2Δ::URA3;lro1Δ::TRP1* | This study |
| *are1Δ/are2Δ/dga1Δ* | *Isogenic to* Scy62 *are1Δ::HIS3;are2Δ::URA3 ;dga1Δ::LEU* | This study |
| *are1Δ/lro1Δ/dga1Δ* | *Isogenic to* Scy62 *are1Δ::HIS3;lro1Δ::TRP1;dga1Δ::URA3* | This study |
| *are2Δ/lro1Δ/dga1Δ* | *Isogenic to* Scy62 *are2Δ::HIS3;lro1Δ::TRP1;dga1Δ::URA3* | This study |

**Supplementary table 4: List of plasmids**

| Serial. No. | Name of the plasmid used | Selectable marker | Source/Reference |
|---|---|---|---|
| 1 | CuGFP-ATG8(416) | Ura | Add gene |
